# Multistep mechanism of DNA replication-coupled G-quadruplex resolution

**DOI:** 10.1101/2020.11.11.378067

**Authors:** Koichi Sato, Nerea Martin-Pintado, Harm Post, Maarten Altelaar, Puck Knipscheer

## Abstract

G-quadruplex (or G4) structures are non-canonical DNA structures that form in guanine-rich sequences and threaten genome stability when not properly resolved. G4 unwinding occurs during S phase via an unknown mechanism. Using *Xenopus* egg extracts, we define a three-step G4 unwinding mechanism that is coupled to DNA replication. First, the replicative helicase (CMG) stalls at a leading strand G4 structure. Second, the DHX36 helicase mediates the bypass of the CMG past the intact G4 structure, which allows approach of the leading strand to the G4. Third, G4 structure unwinding by the FANCJ helicase enables the DNA polymerase to synthesize past the G4 motif. A G4 on the lagging strand template does not stall CMG, but still requires DNA replication for unwinding. DHX36 and FANCJ have partially redundant roles, conferring robustness to this pathway. Our data reveal a novel genome maintenance pathway that promotes faithful G4 replication thereby avoiding genome instability.

## Introduction

Guanine-rich nucleic acid sequences can adopt four-stranded structures, termed G-quadruplexes (G4s) (Gellert et al., 1962). G4s comprise three or more stacked G-quartets, planar structures formed by four guanines connected via Hoogsteen hydrogen bonding and stabilized by a monovalent cation (Sen and Gilbert, 1988; Sundquist and Klug, 1989). Vertebrate genomes contain thousands of sequences that can form G4 structures with a wide variety of topologies, arising from parallel and anti-parallel G-strand direction and intervening loops (Burge et al., 2006). Moreover, these motifs preferentially localize to evolutionarily-conserved regulatory loci (Chambers et al., 2015; Huppert and Balasubramanian, 2005). While G4s are involved in a broad range of biological processes such as transcriptional regulation, telomere maintenance, and epigenetic regulation (Sarkies et al., 2010; Siddiqui-Jain et al., 2002; Zahler et al., 1991), these structures have also been linked to genome instability (Cheung et al., 2002).

The eukaryotic genome encodes at least ten DNA helicases that show G4 unwinding activity *in vitro* (Lerner and Sale, 2019). Furthermore, genetic studies have provided important clues that link several helicases to G4 unwinding *in vivo.* In yeast, the 5’-3’ DNA helicase PIF1 is essential for genome stability at G4s (Myung et al., 2001; Ribeyre et al., 2009). PIF1 is highly conserved in mammals, however its function appears to predominate in the mitochondria (Bannwarth et al., 2016; Snow et al., 2007). In human cells, the 5’-3’ DNA helicase FANCJ plays a critical role in preventing large deletions near G4 motifs (London et al., 2008), although it seems to be dispensable for this role in mice (Matsuzaki et al., 2015). Biallelic mutations in *FANCJ* causes Fanconi anemia (FA), a human cancer predisposition syndrome (Seal et al., 2006). While FANCJ acts in the ‘FA pathway’ that repairs DNA interstrand crosslinks (Ishiai et al., 2017; Niraj et al., 2019), this role appears to be independent of its poorly characterized function in G4 unwinding (Castillo Bosch et al., 2014; Wu et al., 2008). Another 5’-3’ DNA helicase, RTEL1, specifically suppresses telomeric G4 instability (Ding et al., 2004; Sfeir et al., 2009). Finally, DHX36 is a 3’-5’ helicase that exhibits an exceptional affinity for DNA G4 structures *in vitro* (Giri et al., 2011), but its biological function has been studied mostly in the context of RNA G4 processing (Lattmann et al., 2011; Nie et al., 2015; Sauer et al., 2019). This helicase is essential for normal embryogenesis in mice (Lai et al., 2012), yet its role in DNA G4 unwinding *in vivo* remains unknown. Notably, depletion of certain helicases, or addition of G4 stabilizing ligands that inhibit G4 unwinding, induce DNA double strand breaks (DSBs) and can cause chromosomal aberrations (Bharti et al., 2013; London et al., 2008; Rodriguez et al., 2012; Uringa et al., 2012). Altogether, this indicates that G4 structures need to be actively unwound by DNA helicases to maintain genome integrity.

Several lines of evidence suggest that G4 structures are predominantly resolved during DNA replication. First, G4 formation is enhanced in S phase, but quickly reduced upon progression to G2 phase (Biffi et al., 2013; Liu et al., 2016). Second, G4 stabilizing ligands arrest cells in G2 phase with 4N DNA content (Izbicka et al., 1999). Third, DNA damage induced by G4 ligands is replication-dependent (Xu et al., 2017). Fourth, several G4 unwinding helicases, such as PIF1, FANCJ, and RTEL1 associate with the replisome, prevent DNA polymerase from stalling at G4s, and preserve replication fork speed through G4s (Alabert et al., 2014; Castillo Bosch et al., 2014; Dahan et al., 2018; Schwab et al., 2013; Vannier et al., 2013). Fifth, FANCJ-deficient *C.elegans* shows a G4-specific deletion signature that implies replication fork blockage as a cause of these deletions (Cheung et al., 2002; Kruisselbrink et al., 2008; Lemmens et al., 2015). These insights suggest that G4 unwinding is coupled to DNA replication to prevent replisome arrest at the G4 structure. However, direct evidence that a G4 structure acts as a roadblock for the replisome is hitherto missing, and moreover, virtually nothing is known about the mechanism of G4 unfolding during DNA replication.

Here, we used *Xenopus laevis* egg extracts to recapitulate unwinding and replication of defined G4 structures. We identify DHX36 and FANCJ as essential helicases involved in this process. Replication past a G4 structure on the leading strand involves stalling of the replicative polymerase CMG, followed by DHX36-mediated CMG bypass of the intact G4 structure allowing the polymerase to approach the G4, and finally the unwinding of the G4 structure by FANCJ enabling G4 motif replication. Although lagging strand G4 resolution does not involve CMG bypass, it still requires DNA replication and both DHX36 and FANCJ. Our data reveal a replication-dependent and robust mechanism that promotes faithful replication of abundant G4 structures and thereby avoids double-strand DNA breaks and chromosomal instability.

## Results

### FANCJ and DHX36 are required for efficient G4 structure unwinding

Vertebrates contain at least ten conserved DNA helicases that are able to unwind G4 structures *in vitro* (FANCJ, BLM, DDX11, DHX9, DHX36, DNA2, PIF1, RTEL1, XPD, and WRN)(Lerner and Sale, 2019), but whether and how these proteins act under physiological conditions is currently unknown. We previously showed that FANCJ promotes the unwinding of G4 structures during replication of single stranded DNA (ssDNA) templates in *Xenopus* egg extract (Castillo Bosch et al., 2014). However, FANCJ depletion only affected the replication of a subset of these structures, indicating that other G4-unwinding helicases are active in the extract. To identify such helicases, we analyzed the proteins recruited to G4-containing plasmids by mass spectrometry. To this end, ssDNA plasmids containing a defined parallel G4 or a control sequence (pBS-G4 and pBS-CON; Figure S1A) were replicated in a high-speed supernatant (HSS) of total *Xenopus* egg lysate. Plasmids were pulled down using beads conjugated with LacI, at times when the polymerase is stalled at the G4 (1.5 and 4.5 minutes), or when the plasmid is fully replicated (50 minutes) (Figure 1A and S1B). Mass spectrometry detected eight G4-unwinding helicases on replicating plasmids (Figure 1B). Some of them were detected on both pBS-G4 and pBS-CON, while others, including FANCJ and DHX36, were specifically enriched on pBS-G4. Moreover, addition of the G4 stabilizing ligand PhenDC_3_ further increased the enrichment of several helicases. The most strongly enriched helicase was DHX36, which peaked at the earliest time point. To examine the role of DHX36 in DNA G4 unwinding, we immunodepleted it from extract and analyzed replication products of G4 templates by denaturing electrophoresis (Figure 1C and D). Depletion of DHX36 initially enhanced polymerase stalling compared to the mock-depleted extract, but all G4 substrate molecules were eventually replicated (Figure 1D). In contrast, FANCJ depletion showed extended polymerase stalling on a subset of the template molecules as shown previously (Figure 1D) (Castillo Bosch et al., 2014). This suggests that DHX36 and FANCJ have different roles in G4 replication. Remarkably, when we depleted both DHX36 and FANCJ, extensive replication stalling at the G4 was observed with almost no replication past the structure for at least an hour (Figure 1D). Addition of recombinant wild-type DHX36 or FANCJ (Figure S1C) restored replication to the respective single depletion levels (Figure 1D). These data indicate that both DHX36 and FANCJ are required for efficient unwinding of G4 structures and that they act partially redundantly.

**Figure 1.**
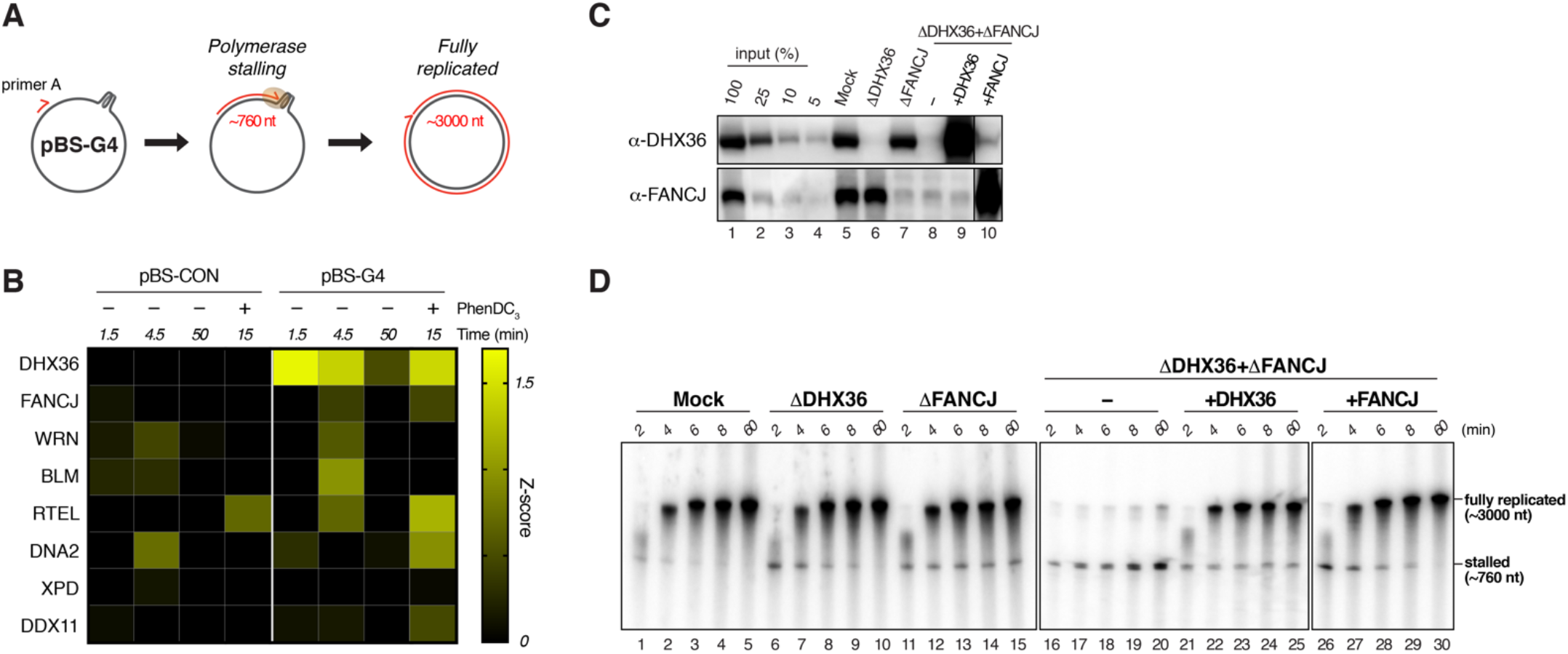
G4 unwinding during replication of ssDNA templates requires DHX36 and FANCJ. (A) Schematic representation of pBS-G4 replication in HSS. Replication is initiated from a primer annealed ~760 nucleotides (nt) upstream of a G4. See also Figure S1A and S1B. (B) pBS-G4 or pBS-CON were replicated in HSS and isolated at various times by LacI pull-down (Budzowska et al., 2015; Larsen et al., 2019). Proteins bound to the plasmids were identified by mass spectrometry (MS) analysis. Relative abundance of proteins is represented by a heatmap showing the mean of the Z scores from four biological replicates. (C) Mock-depleted, DHX36-depleted, FANCJ-depleted, and DHX36-FANCJ-depleted HSS supplemented with buffer, wild-type (WT) FANCJ, or WT DHX36 were analyzed by Western blot with DHX36 and FANCJ antibodies, alongside a dilution series of undepleted extract. (D) pBS-G4 was replicated in the extracts described in c) in the presence of ^32^P-a-dCTP. Replication products were extracted, separated by denaturing PAGE, and visualized by autoradiography. Nascent strands stalled at the G4 sequence (~700 nt) and fully replicated molecules (~3000 nt) are indicated.

### G4 structure unwinding and replication on dsDNA templates

Replication of ssDNA substrates in *Xenopus* egg extract does not involve the complete replisome because unwinding of the DNA duplex is not required. To investigate the role of DHX36 and FANCJ at *bone fide* replication forks, we set out to develop a system that would recapitulate G4 unwinding and replication in the context of a double stranded DNA (dsDNA) template. Replication of dsDNA in extract involves incubation in HSS to allow the assembly of prereplication complexes, followed by addition of a concentrated nucleoplasmic extract (NPE) to promote initiation and a single round of DNA replication. We constructed dsDNA plasmids that contain a G4 motif or a control sequence similar to the ssDNA templates (pdsG4^BOT^ and pdsCON; Figure S2A and B). To monitor replication intermediates, we separated replication products on a denaturing agarose gel after linearization with HincII. Replication of pdsG4^BOT^ and pdsCON quickly yielded full length molecules and no intermediates accumulated (Figure S2C). This suggests that a stable G4 structure does not form in this context, which is consistent with duplex DNA hydrogen bonds being more stable than G4 hydrogen bonds (Phan and Mergny, 2002).

We then prepared dsDNA plasmids containing a short non-complementary region, in which the G4 structure can be pre-formed on the top or bottom strand (pG4^TOP^ and pG4^BOT^), along with non-G4 control plasmids (pCON and pPolyT; Figure S2A). Replication of control plasmids yielded full-length products without accumulation of any intermediates, indicating that the non-complementary region does not hinder the replication machinery (Figure 2A). However, replication of pG4^BOT^ and pG4^TOP^ resulted in a transient accumulation of ~2.0 kb and ~3.6 kb products that were quickly converted to full-length products (Figure 2A). The accumulation of these fragments was greatly enhanced by PhenDC_3_ (Figure 2B) and suppressed when the G4 structure was not pre-formed by omitting potassium (Figure S2D). These data suggest that replication temporarily halts at the site of the G4 structure and resumes upon G4 unwinding. Notably, the 2.0 kb fragment is less defined and prone to degradation in pG4^TOP^, similar to the 3.6 kb fragment in pG4^BOT^, suggesting that these are lagging strand products with free 5’ ends (Figure S2E). This indicates that replication stalling occurs on the G4 structure-containing strand, which was confirmed by digestion with ClaI, which only cleaves the replicated top strand (Figure S2A and F).

**Figure 2.**
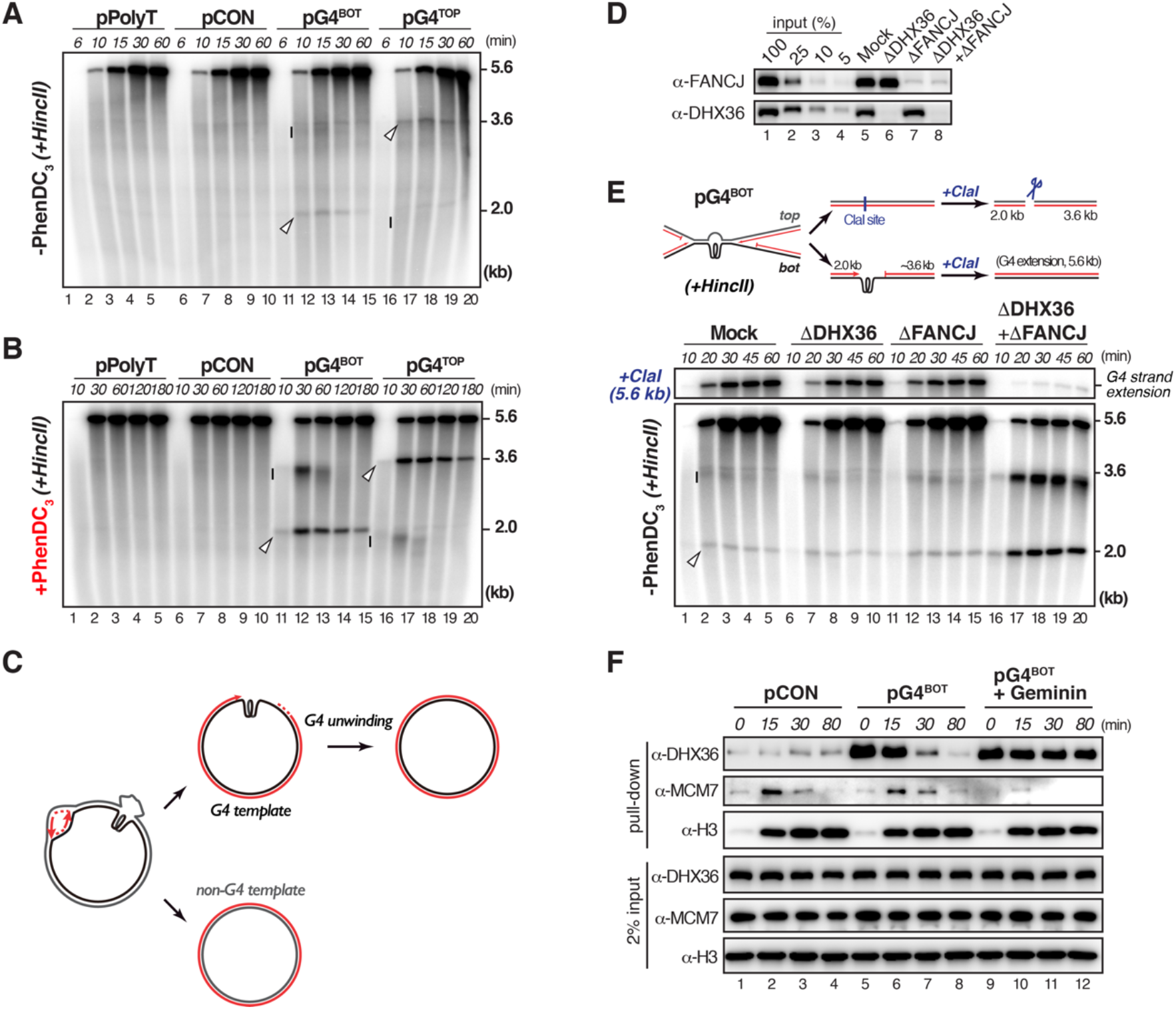
DHX36 and FANCJ are essential for G4 unwinding on physiological replication forks. (A and B) pPolyT, pG4^MUT^, pG4^BOT^ and pG4^TOP^ were replicated without (A) or with PhenDC_3_ (B) in the presence of a-^32^P-dCTP. Replication intermediates were linearized with HincII and analyzed by denaturing agarose gel electrophoresis and autoradiography. 2.0 kb and ~3.6 kb nascent products observed during pG4^BOT^ replication are indicated with an arrowhead and a bar, respectively. For pG4^TOP^ ~2.0 and 3.6 kb products are indicated with a bar and an arrowhead, respectively. (C) Model for pG4^BOT^ replication. Replication of the G4 template strand involves polymerase stalling, while replication of the non-G4 template proceeds without stalling. (D) Mock-depleted, DHX36-depleted, FANCJ-depleted, and DHX36-FANCJ-depleted NPE were analyzed by Western blot with DHX36 and FANCJ antibodies alongside a dilution series of undepleted extract. (E) pG4^BOT^ was replicated in the indicated extracts in the presence of a-^32^P-dCTP. Replication intermediates were digested with HincII (lower autoradiogram), or HincII and ClaI (upper autoradiogram) and analyzed as in (A). 2.0 kb and ~3.6 kb digestion products are indicated with an arrowhead and a bar, respectively. Since ClaI specifically cuts the replicated top strand, the 5.6 kb full length product results only from the replicated bottom strand that contains the G4 (schematic, top panel). See also Figure S2F. (F) pCON and pG4^BOT^ plasmids were replicated in *Xenopus* egg extract. At various time points (t=0 was collected immediately after NPE addition) plasmids were isolated by LacI pull-down. Proteins bound to the plasmids were analyzed, along with a 2% input sample, by Western blot analysis with the indicated antibodies. Where indicated, HSS was supplemented with Geminin to inhibit DNA replication initiation.

Even in the presence of PhenDC_3_, when replication stalling was prominent, extension products still readily appeared (Figure 2B). This suggests that replication of the non-G4 strand precedes G4 structure unwinding. In agreement with this, strand specific digestion by ClaI showed that replication of non-G4 strand was not significantly affected by PhenDC_3_ (Figure S2F). Furthermore, two-dimensional gel electrophoresis demonstrated that the sister chromatids were efficiently resolved upon replication fork stalling at the G4 (Figure S2G). Therefore, we conclude that replication of G4-containing dsDNA templates results in transient replication stalling on the G4 strand, while replication of the opposite strand is not interrupted (Figure 2C).

### DHX36 and FANCJ promote replication-dependent G4 unwinding

We next explored the role of DHX36 and FANCJ in this system. Similar to what we observed for ssDNA substrates, double depletion of DHX36 and FANCJ greatly increased replication stalling at the G4 structure and severely compromised extension of the G4 strand, while each single depletion had only a minor effect (Figure 2D and E). This indicates that replication of the G4 sequence requires G4 unwinding by DHX36 and FANCJ. This unwinding could occur before or during DNA replication. To examine this, we pulled down replicating plasmids and monitored DHX36 binding. DHX36 was detected specifically on G4-containing plasmids prior to replication initiation and was displaced over time, indicative of G4 unwinding during replication (Figure 2F). Importantly, when replication initiation was abrogated by addition of Geminin, DHX36 persisted on pG4^BOT^ (Figure 2F). This indicates that DNA replication promotes G4 structure resolution. Early accumulation of DHX36 at the G4 was also observed in our mass spectrometry data in which FANCJ peaked later (Figure 1B), suggesting that DHX36 and FANCJ act at different stages of G4 unwinding.

### G4 structures are resolved during leading and lagging strand synthesis

Since replication forks in our system arrive at the G4 structure either from the left or the right, and fork convergence occurs rapidly on small plasmids, we cannot distinguish whether the G4 is resolved during leading or lagging strand synthesis. To allow this distinction, we made use of an array of lac operator (*lacO*) repeats situated downstream of the G4 structure (Figure S2A). Upon pre-incubation with the Lac repressor (LacI), arrival of the leftward fork is efficiently blocked (Figure S3A and B) and only the rightward fork encounters the G4 on the leading (in pG4^BOT^) or lagging strand (in pG4^TOP^) (Figure S3C-D, cartoons). This also more closely resembles the *in vivo* situation, since larger inter-origin distance will make fork convergence less likely (Berezney et al., 2000). We replicated pG4^TOP^ and pG4^BOT^ in the presence of LacI (and under these conditions will refer to them as pG4^Lag^ and pG4^Lead^, respectively). Analysis of the digested replication products on a sequencing gel showed that the leading and lagging strands were rapidly extended past the G4^Lead^ and G4^Lag^ respectively (Figure S3C and D). The extension kinetics were comparable to that of pCON and a convergent fork situation. Therefore, both leading and lagging strand G4s are efficiently resolved, and dual fork collision is not required for G4 structure unwinding.

### Leading strand G4 replication is a three-step process

To examine how a G4 structure on the leading strand is resolved, we further analyzed the replication products of pG4^Lead^. Two faint nascent strand clusters appeared within 15 minutes and declined to undetectable levels by 30 minutes (Figure S3C, blue and green brackets). G4 stabilization by PhenDC_3_ enhanced these clusters that are formed by initial stalling of the leading strand 13 to 26 nt upstream of G4^Lead^ (‘-13 to −26’ products), followed by a second stalling event at 1 to 3 nt from the G4^Lead^ (‘-1 to −3’ products) (Figure 3A and B). DNA lesions such as ICLs and DNA-protein crosslinks (DPCs) arrest leading strand synthesis roughly 20 to 30 nt from the damage-site due to steric hindrance by CMG helicase that translocates along the leading strand template ahead of the polymerase (Duxin et al., 2014; Räschle et al., 2008). Therefore, the −13 to −26 products likely reflect the CMG footprint upon encountering the G4. Consistent with this, when the G4 was present on the lagging strand template, the −13 to −26 products were hardly detected, while the −1 to −3 products still accumulated. Moreover, the −13 to −26 products reappeared when LacI was omitted and both lagging and leading strand synthesis occurred (Figure 3B, lanes 15-21).

**Figure 3.**
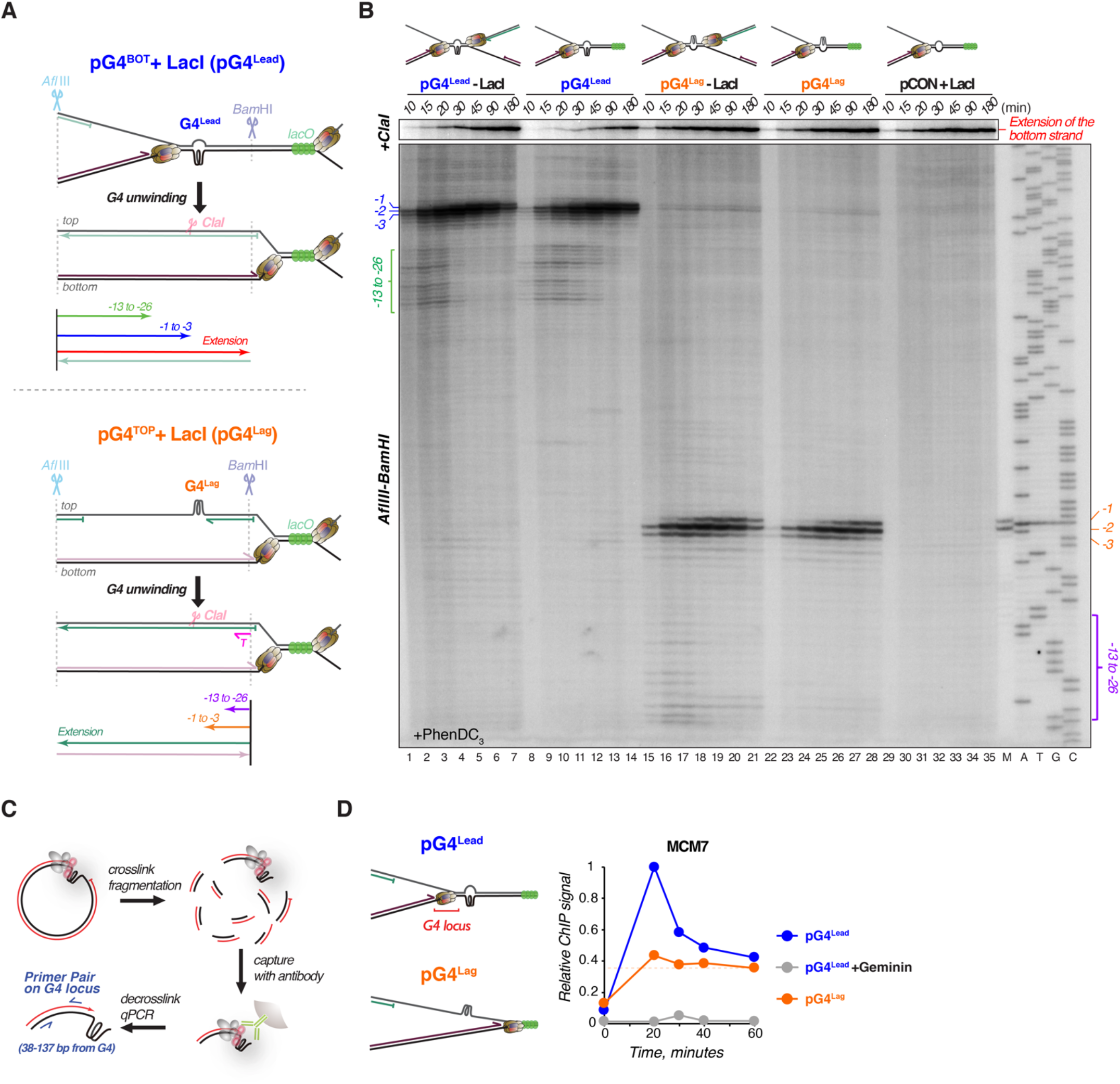
Multistep replication bypass of a G4 structure. (A) Schematic representation of nascent leading and lagging strand products generated during pG4^Lead^ or pG4^Lag^ replication. AflIII and BamHI digestion sites are indicated. Digestion with ClaI specifically digests the extension products of the top strand. Brown hexamer, CMG; Green sphere, LacI; T, sequencing primer T. (B) pG4^Lead^, pG4^Lag^, and LacI-bound pCON were replicated in the presence of PhenDC_3_, digested with AflIII, BamHI and ClaI (upper gel) or AflIII and BamHI (lower gel), and separated on denaturing polyacrylamide gel, and visualized by autoradiography. Where indicated, LacI was omitted. A sequencing ladder derived from extension of primer T annealed to pG4^TOP^, and a 97-bp and 96-bp PCR product (M, see methods) were used as size markers. Nascent strands corresponding to the −13 to −26 products and the −1 to −3 products are indicated with brackets and bars, respectively. (C) Scheme of the chromatin immunoprecipitation (ChIP) assay. G4 containing plasmids were replicated, DNA and proteins (ovals) were crosslinked, samples were sonicated, and immunoprecipitated with the antibody of interest (green). The co-precipitated DNA was amplified by quantitative PCR (qPCR) with primers (blue arrows) specific to the G4 locus. (D) pG4^Lead^ and pG4^Lag^ were replicated and analyzed by ChIP-qPCR with the MCM7 antibody and a primer pair for the G4 locus (schematic, left panel). Where indicated, Geminin was added to inhibit replication initiation. Since DNA fragments analyzed by ChIP are on average ~600 bp long, some DNA fragments will contain both the G4 and *lacO* loci (420 bp apart). This likely generates the background signal on pG4^Lag^ in which MCM7 is stalled at the LacI-bound *lacO* array (dashed line).

To confirm that CMG stalls at a leading strand G4, we monitored the accumulation of MCM7, a CMG subunit, by plasmid-based chromatin immunoprecipitation (ChIP) (Figure 3C). During replication of pG4^Lag^, background levels of MCM7 were detected at the G4 site, most likely due to arrival of CMG at the *lacO* locus (Figure 3D). However, during the replication of pG4^Lead^, MCM7 accumulated at the G4 site, showing a strong peak at 20 minutes (Figure 3D) that coincided with the presence of −13 to −26 products (Figure S3C). This indicates that CMG stalling leads to the accumulation of these leading strand products. All together, these data suggest that replication past a G4 structure on the leading strand is a three-step process (Extended Data Fig. 3e). First, CMG stalls at a G4 structure resulting in the accumulation of leading strand products 13 to 26 nt from the G4. Subsequently, CMG vacates the G4 site, which enables the polymerase to approach the G4 where it stalls again at 1 to 3 nt from the G4. Finally, the G4 structure is unwound to allow replication of the G4 sequence. Because a G4 on the lagging strand is not encountered by CMG, its replication only involves the latter two steps, polymerase stalling at the G4 followed by G4 unwinding and replication (Figure S3E).

### DHX36 and FANCJ collaborate to unwind a leading strand G4

We next examined the roles of FANCJ and DHX36 in these two steps. During pG4^Lead^ replication, depletion of DHX36 resulted in a transient accumulation of the −13 to −26 products, whereas depletion of FANCJ enhanced the −1 to −3 products (Figure 4A, arrowheads). Double depletion of DHX36 and FANCJ strongly enhanced both stalling clusters and severely impaired the extension past the G4 structure (Figure 4A, lanes 16-21). The deficiency was restored to single depletion levels by addition of wild-type DHX36 and FANCJ proteins, but not their ATPase mutants (Figure S1C and S4A-D). Therefore, the major role of DHX36 during leading strand G4 replication is likely to promote polymerase approach by facilitating CMG clearance from the G4 site, while FANCJ mostly acts in G4 unwinding, although they also act in part redundantly in both processes. In agreement with this, depletion of DHX36 had no effect on G4^Lag^ replication that does not involve CMG stalling (Figure S4E). FANCJ depletion caused polymerase stalling on the lagging strand which was further enhanced by FANCJ/DHX36 double depletion (Figure S4E), indicating that DHX36 and FANCJ also act redundantly in G4 unwinding on the lagging strand.

**Figure 4.**
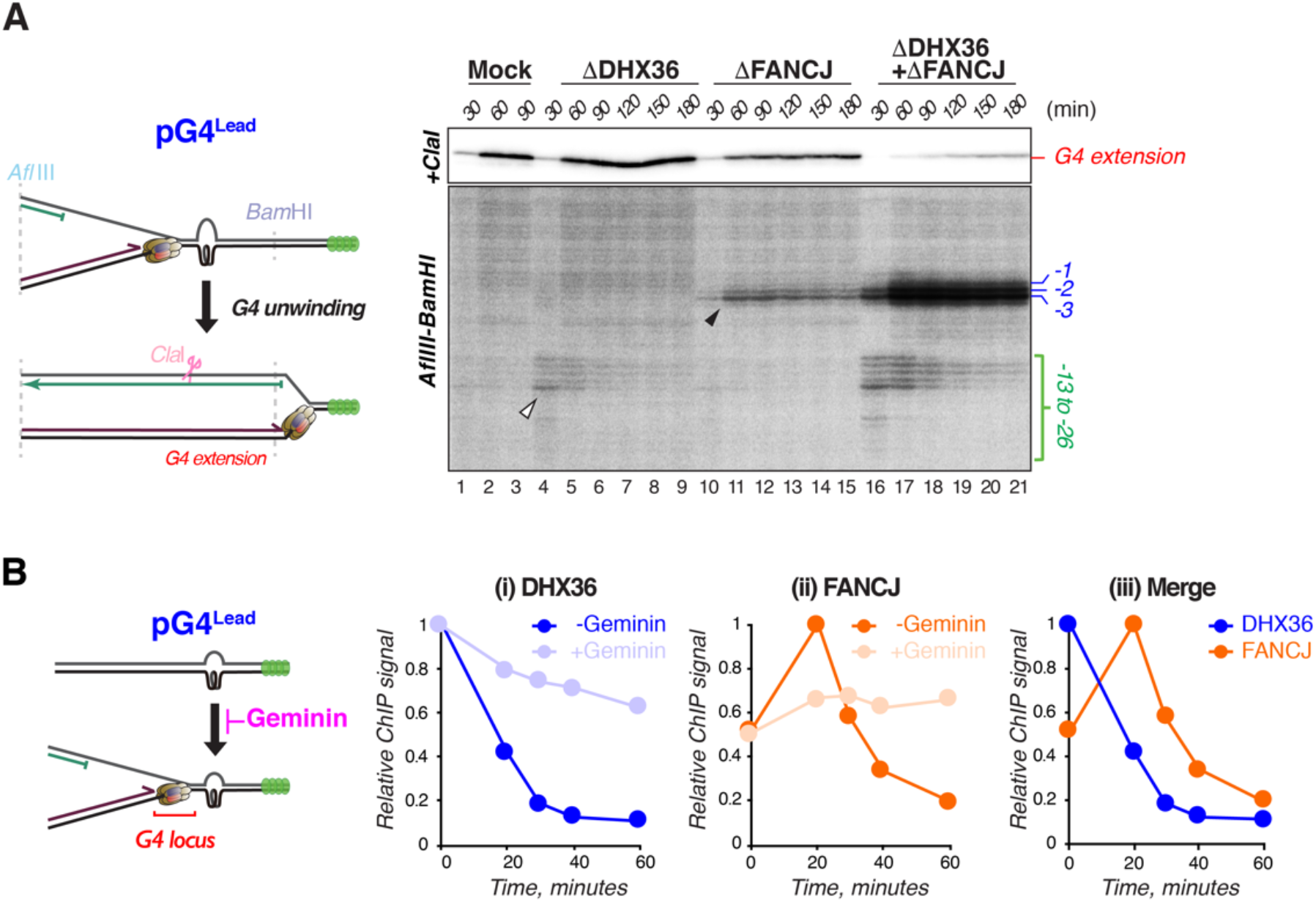
DHX36 and FANCJ cooperate in unwinding a leading strand G4 structure. (A) pG^Lead^ was replicated in the indicated extracts, digested with AflIII, BamHI, and ClaI (upper gel) or with AflIII and BamHI (lower gel), separated on denaturing polyacrylamide gel, and visualized by autoradiography. The −13 to −26 products and the −1 to −3 products are indicated with a bracket and bars, respectively. Schematic representation of the generated fragments is shown on the left. Brown hexamer, CMG; green sphere, LacI. (B) pG4^Lead^ was replicated, and analyzed by ChIP-qPCR with DHX36 (i) and FANCJ (ii) antibodies using a primer pair for the G4 locus (schematic representation in left panel). Where indicated, Geminin was added to inhibit replication initiation. To compare the timing of accumulation for both proteins, DHX36 and FANCJ ChIP data in the absence of Geminin were plotted together in one graph (iii).

To gain insight into how FANCJ and DHX36 collaborate to replicate a G4 on the leading strand template, we monitored their recruitment to G4^Lead^ by ChIP. Consistent with our plasmid pull-down assay, DHX36 was already present at G4^Lead^ prior to replication initiation and mostly disappeared by 30 minutes (Figure 4B, i). In contrast, while some FANCJ was present prior to replication, it further accumulated at G4^Lead^ simultaneously with the disappearance of DHX36, and FANCJ only vacated the G4 site at later times (Figure 4B, ii and iii). Interestingly, inhibition of replication initiation by geminin caused persistent DHX36 binding at the G4^Lead^ and prevented FANCJ from accumulating (Figure 4B). Similar results were obtained with pG4^Lag^ (Figure S4F). These data indicate that DHX36 and FANCJ are sequentially bound to both leading and lagging strand G4s during replication. Since DHX36 binds to the G4 structure prior to FANCJ, and its depletion induced CMG stalling at the G4, we envision that it facilitates CMG clearance to allow polymerase approach to the G4 site.

### DHX36 promotes CMG bypass by generating ssDNA downstream of the G4

The CMG helicase could vacate the G4 site by two distinct mechanisms, unloading or bypass (Fullbright et al., 2016; Sparks et al., 2019; Wu et al., 2019). To distinguish between these possibilities, we replicated pG4^Lag^ and pG4^Lead^, and analyzed the accumulation of MCM7 at the region flanking the *lacO* array by ChIP (Figure 5A). If CMG is unloaded at the leading strand G4, we would expect no MCM7 to be detected at the LacI-bound *lacO* locus for pG4^Lead^, while it would accumulate at the locus in pG4^Lag^. As anticipated, MCM7 was readily detected at the *lacO* locus on pG4^Lag^ in the presence of LacI (Figure 5A). This accumulation was due to the LacI-bound *lacO* array as it was abolished in the absence of LacI. On pG4^Lead^, we observed MCM7 accumulation to similar levels, but with a delay of about 20 minutes (Figure 5A). This delay is consistent with the initial accumulation of MCM7 at the leading strand G4, followed by the loss of MCM7 at the G4 that coincides with accumulation at the *LacO* locus (Figure 3D and Figure 5A). Therefore, we conclude that DHX36 facilitates CMG bypass of the G4^Lead^, rather than extracting it from the DNA. Consistent with this, inhibition of the CMG unloader p97 (Fullbright et al., 2016; Semlow et al., 2016) did not enhance the CMG footprint at the leading strand G4 (Figure S5A-C).

**Figure 5.**
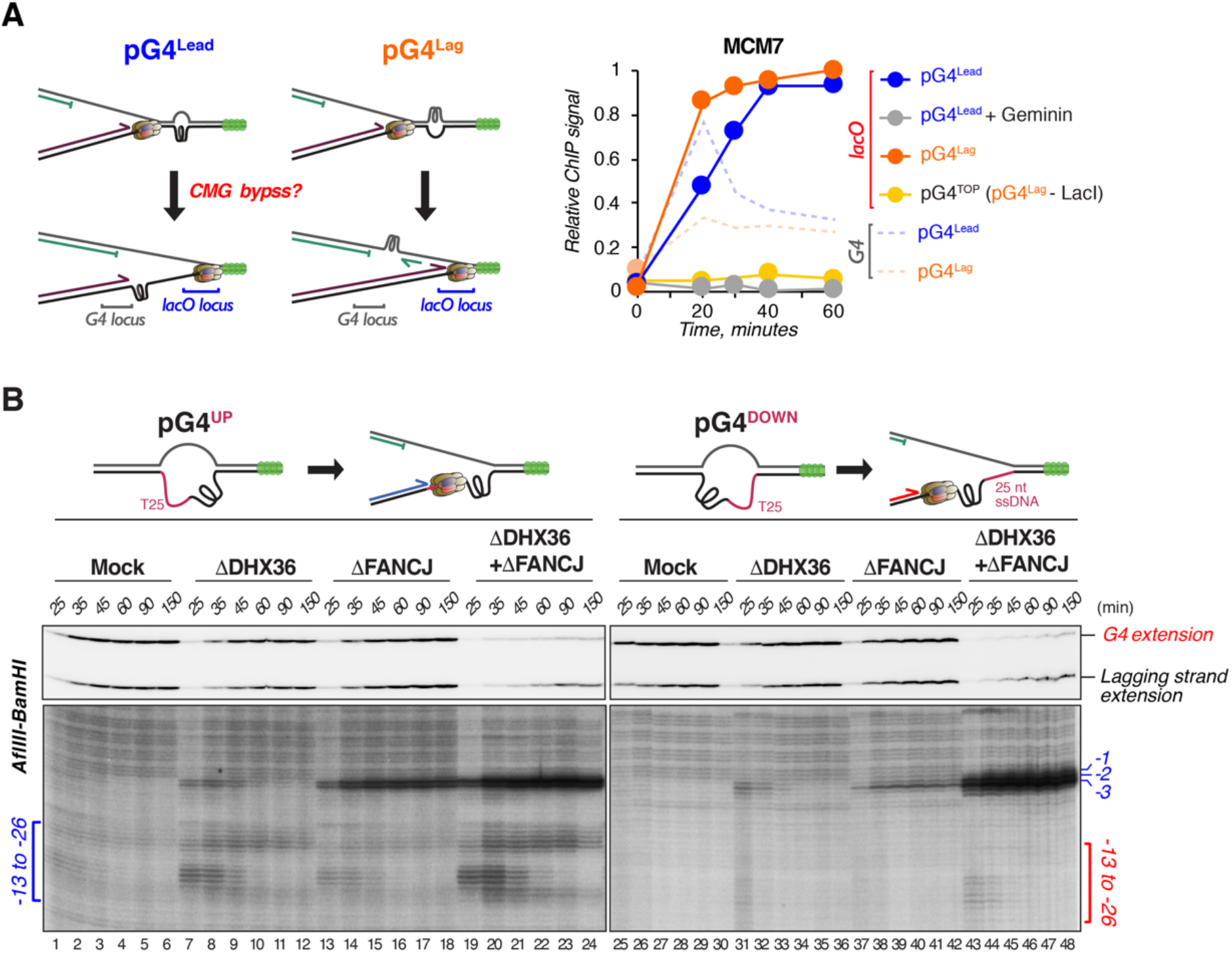
DHX36 promotes CMG bypass by generating ssDNA downstream of the G4 structure. (A) pG4^Lead^ and pG4^Lag^ were replicated, and analyzed by ChIP-qPCR with the MCM7 antibody using a primer pair for the *lacO* locus (295-388 bp downstream from the G4) or the G4 locus. Where indicated, LacI was omitted, or Geminin was added to inhibit replication initiation. The MCM7 levels on the G4 locus (from Figure 3D) were added to the graph for comparison. If CMG bypasses the G4, it will be detected on the *lacO* locus as depicted in the left panel. (B) pG4^UP^ and pG4^DOWN^ were replicated, digested with AflIII and BamHI, separated on denaturing polyacrylamide gel, and visualized by autoradiography. The −13 to −26 products and the −1 to −3 products are indicated with brackets and bars, respectively.

Since a G4 recognition mutant of DHX36 (DHX36^Y53A^) did not promote CMG bypass (Figure S4A and B), G4 binding of DHX36 is critical for efficient bypass. In addition, DHX36 is reported to translocate and unwind duplex DNA with a 3’-5’ directionality (Yangyuoru et al., 2018), and could therefore generate ssDNA downstream of the G4 structure. Generation of ssDNA by helicase mediated unwinding has recently been shown to promote CMG bypass of a DNA-protein crosslink (Sparks et al., 2019). We therefore examined whether downstream ssDNA is sufficient for the CMG to bypass the G4. To this end, we inserted a non-complementary 25 nt polyT sequence downstream of the leading strand G4 (pG4^DOWN^; Figure 5B). As a control, we also prepared a plasmid in which the polyT stretch was inserted upstream of the G4 (pG4^UP^). While replication of pG4^UP^ resulted in the accumulation of CMG stalling products upon DHX36 and DHX36-FANCJ depletion, these were largely absent when pG4^DOWN^ was replicated (Figure 5B). This indicates that ssDNA downstream of the G4 structure is sufficient for CMG bypass. Consistent with this, CMG stalling at a G4 in a DHX36/FANCJ-depleted extract was rescued by fork convergence (Figure S5D). However, while G4 bypass was facilitated by arrival of the second fork or by downstream ssDNA in pG4^DOWN^, DNA synthesis past the G4 was still potently blocked (Figure 5B), indicating that the CMG bypasses the intact G4 structure (Figure S5E).

### CMG bypass is essential for FANCJ-mediated G4 unwinding on the leading strand

Given the role of DHX36 in CMG bypass, and our observation that FANCJ replaces DHX36 at the G4 structure during G4 resolution, we hypothesized that CMG bypass might be a prerequisite for this helicase switch. To test this, we purified an ATPase mutant of DHX36 which is defective in translocation (DHX36^E327A^, Figure S1C) (Tran et al., 2004), and first examined its localization to the G4 during replication by ChIP. Consistent with a defect in translocation, we found this DHX36^E327A^ mutant to be retained at the G4 site in contrast to the wildtype protein that readily vacates the site during replication (Figure 6A and B, i). The DHX36^E327A^ mutant also impaired CMG bypass as seen by the enhanced levels of MCM7 at G4^Lead^ locus and the reduced MCM7 levels at the *lacO* locus (Figure 6B, ii and iv). Moreover, the DHX36^E327A^ mutant prevented accumulation of FANCJ at the G4^Lead^ during replication (Figure 6B, iii). Consistent with a lack of FANCJ accumulation, the mutant enhanced the CMG stalling signature and extended it to at least 2.5 hours (Figure 6C), indicating that the G4 structure is hardly unwound. Conversely, when CMG bypass was facilitated by allowing replication fork convergence, FANCJ accumulation was restored (Figure S6A) and the replication defect of G4^Lead^ in the presence of the DHX36 mutant was rescued (Figure 6D). Taken together, these data indicate that CMG bypass is required for G4 unwinding most likely because it promotes FANCJ accumulation at the leading strand G4.

**Figure 6.**
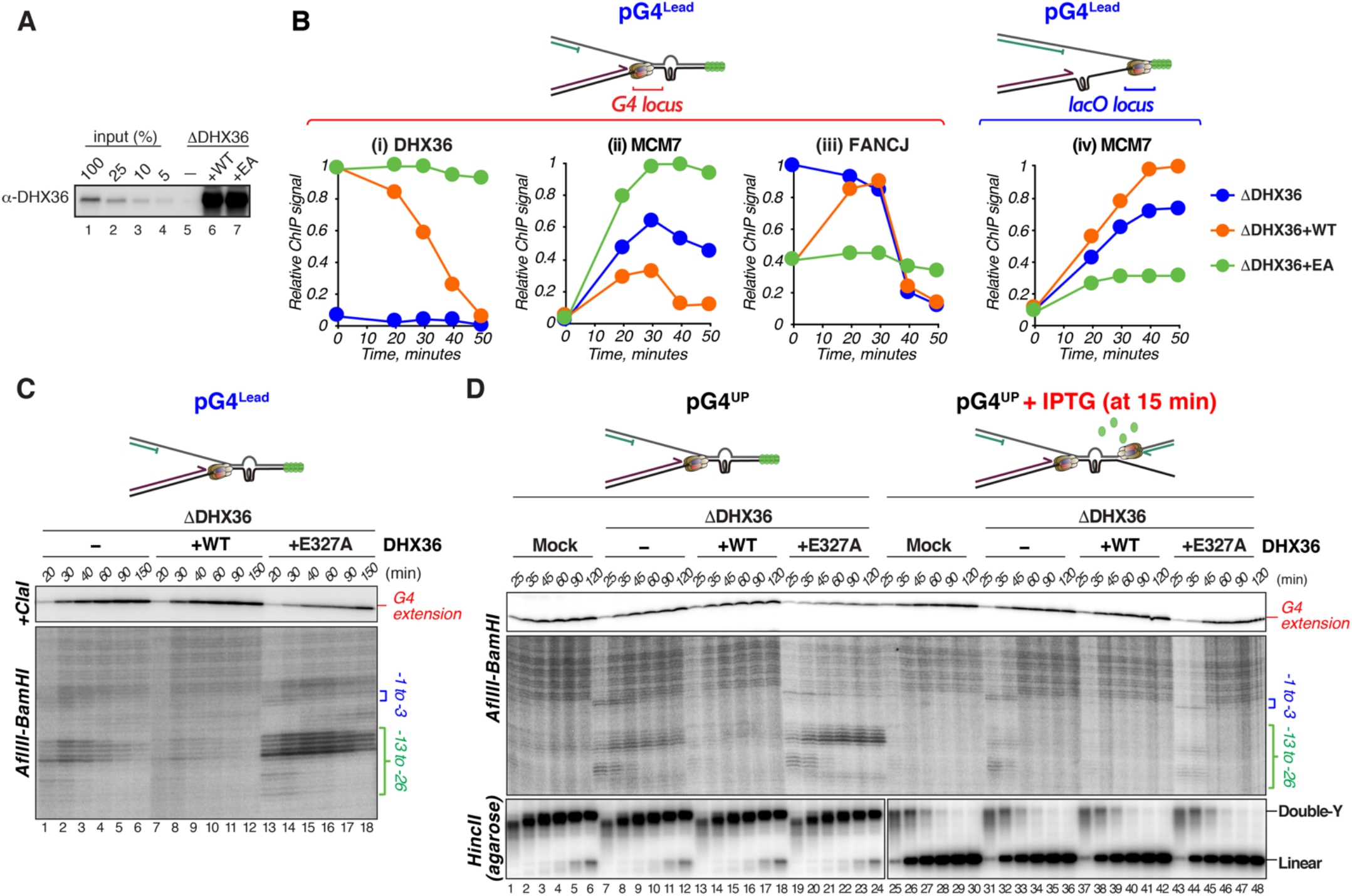
CMG bypass is a pre-requisite for unwinding of a leading strand G4. (A) DHX36-depleted NPE supplemented with buffer, wild-type (WT) DHX36, or DHX36^E327A^ (EA) were analyzed by Western blot with the DHX36 antibody alongside a dilution series of undepleted extract. (B) pG4^Lead^ was replicated in the indicated extracts, and analyzed by ChIP-qPCR with DHX36 (i), MCM7 (ii and iv), or FANCJ (iii) antibodies using a primer pair for the G4 locus (i-iii), or for the *lacO* locus (iv). Schematic representation of qPCR loci on top. Brown hexamer, CMG; green sphere, LacI. (C) pG4^Lead^ was replicated in the indicated extracts, digested with AflIII, BamHI, and ClaI (upper gel), or AflIII and BamHI (lower gel), separated on a denaturing polyacrylamide gel, and visualized by autoradiography. The −13 to −26 and −1 to −3 products are indicated with brackets. (D) pG4^UP^ (G4 on bottom strand and T25 upstream of G4) was replicated in the indicated extracts, and analyzed as in (C) but only digested with AflIII and BamHI. Extension product of the nascent leading strand past the G4 is shown in the upper panel. Where indicated, LacI was released by addition of IPTG at 15 minutes (lanes 25-48). The −13 to −26 and −1 to −3 products are indicated with brackets. To monitor replication fork block by LacI, extracted samples were digested with HincII, separated on native agarose gel, and visualized by autoradiography (bottom gels).

### DHX36 and FANCJ have partially redundant roles

Since the CMG stalling signature is more extensive in the FANCJ-DHX36-depleted extract compared to the DHX36-depleted extract (Figure 4A), it seems that FANCJ can partially replace the function of DHX36 in CMG bypass. Consistent with this, FANCJ levels at the G4 region are enhanced prior to replication initiation in a DHX36-depleted extract (Figure S6A, ii). Since FANCJ is a 5’-3’ DNA helicase, it could translocate along the non-G4 strand to generate ssDNA downstream of the G4 structure to facilitate CMG bypass. To test this possibility, we introduced a stable DNA-protein crosslink on the non-G4 strand of pG4^Lead^ to prevent FANCJ translocation (pG4^Lead-DPC^, Figure S6B) (Duxin et al., 2014). During pG4^Lead-DPC^ replication, the leading strand mostly extended past the G4 as observed for a control plasmid that did not contain DPC (pG4^Lead-CON^; Figure S6C and D), indicating that the DPC does not affect the G4^Lead^ replication. Strikingly, the CMG stalling footprint (−13 to −26 cluster) in the DHX36-depleted extract was enhanced by the presence of the DPC to the same extend as observed in a DHX36-FANCJ double depleted extract (Figure S6D). These data indicate that, in absence of DHX36, FANCJ facilitates CMG bypass by translocating on the non-G4 strand. DHX36 can also partially replace FANCJ’s function in G4 unwinding (Figure 1D and 4A). Consistent with this, DHX36 appears to persist longer on both leading and lagging strand G4s in the absence of FANCJ (Figure S6E). Collectively, DHX36 and FANCJ have partially redundant roles during both CMG bypass and G4 unwinding processes.

## Discussion

This work establishes the first comprehensive mechanism of how G4 structures are resolved during DNA replication. On the leading strand template, this requires three steps including CMG bypass of the intact G4 structure, while on the lagging strand only the final two steps are required, polymerase stalling at the G4 and unwinding of the G4 structure (Figure 7). CMG bypass is initiated by collision of the CMG helicase to the G4 structure that promotes ssDNA generation by DHX36. This minimizes the time required for CMG bypass while avoiding unscheduled ssDNA generation which could induce DSBs. In addition, on both strands, approach of the DNA polymerase to the G4 is required for the helicase switch from DHX36 to FANCJ and subsequent unwinding of the structure (see further discussion below). This ensures tight coupling between G4 unwinding and synthesis past the G4, which avoids reformation of the structure between these events. Partial redundancy between DHX36 and FANCJ make this mechanism further robust (Figure S7A).

**Figure 7.**
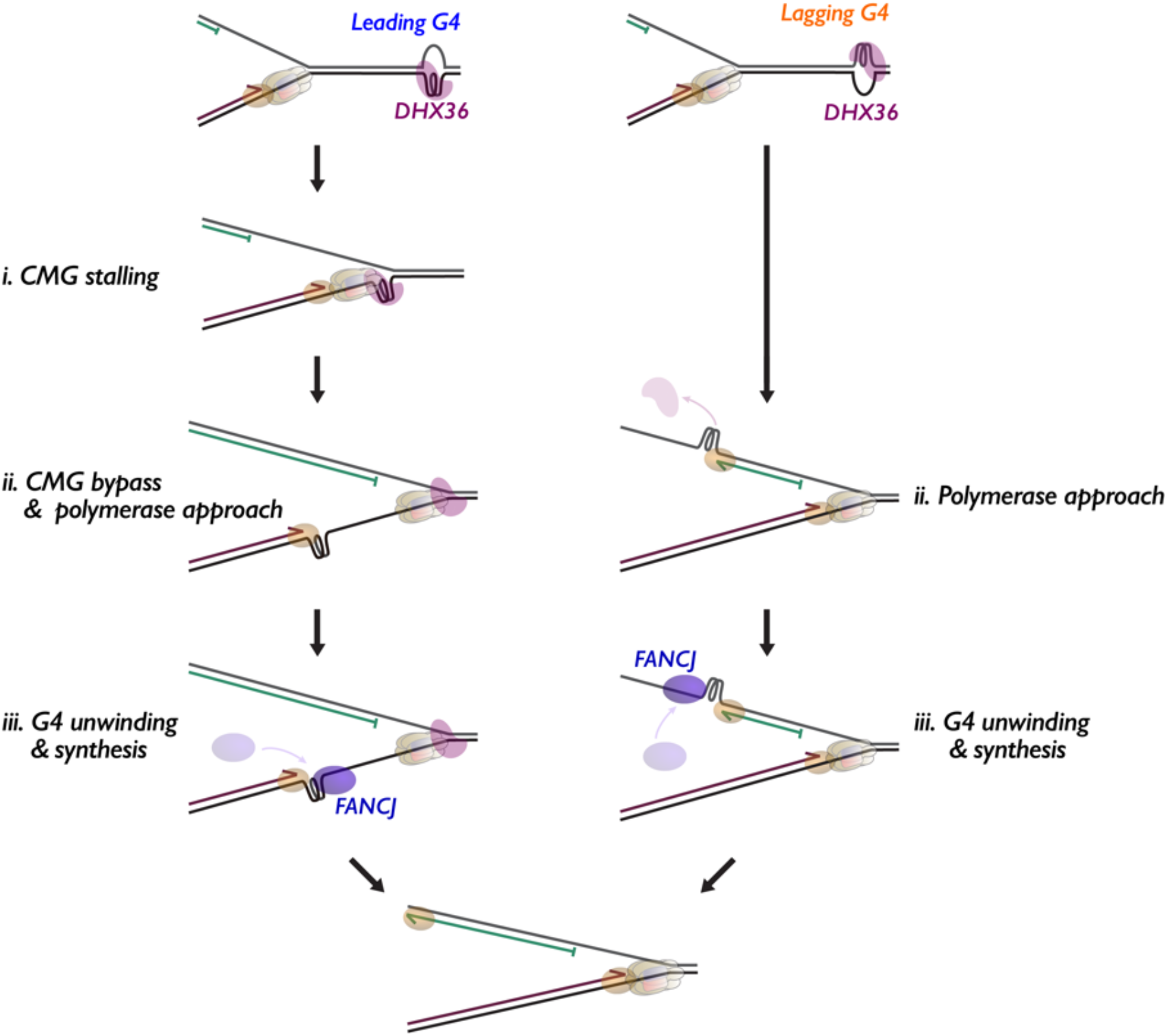
Model for replication-coupled G4 structure unwinding. Models for replication of a leading strand G4 (left) and a lagging strand G4 (right). See Discussion for details. Brown hexamer, CMG; magenta oval, DHX36; blue oval, FANCJ; brown oval, the leading strand DNA polymerase; green oval, the lagging strand DNA polymerase.

### Mechanism of CMG stalling at a leading strand G4

During leading strand synthesis, the CMG helicase is the first replisome component that encounters the G4 structure. This causes CMG to stall and induces a leading strand stalling footprint 13 to 26 nt upstream of the G4 structure (Figure S3C), which is enhanced in the absence of FANCJ and DHX36 (Figure 4A, lanes 16-21). The −21 to −26 cluster is relatively faint and quickly converted to another, −13 to −17, cluster. These observations suggest that CMG stalls at the G4 in two steps. The initial −21 to −26 cluster likely reflects stalling of the intact CMG at the G4 structure (Figure S7B, i), because a similar footprint was observed at an interstrand crosslink and a DPC (Duxin et al., 2014; Räschle et al., 2008), and this is consistent with the CMG structure. MCM2-7, the hexameric subunit of the CMG that encircles the DNA, consists of a distinct N-terminal and C-terminal tier (Li et al., 2015) and translocates on the leading strand template with the N-terminal tier in front (Georgescu et al., 2017). The N-tier channel has a ~20-Å-diameter opening (Li et al., 2015), but this does not fit a G4 with a minimal diameter of ~24 Å (Figure S2B) (Do and Phan, 2012). While this explains the first CMG arrest, the second arrest, 13 to 17 nt from the G4 structure, raises the interesting possibility that under certain conditions the G4 can be accommodated within the N-tier channel (Figure S7B, ii). We envision at least two mechanisms that could support this. First, CMG changes its conformation upon stalling to create space for the G4 structure. In support of this model, purified MCM complex was shown to accommodate a ~5 kDa DPC adduct (Nakano et al., 2013). Alternatively, CMG or another DNA helicase might remodel the G4 structure to allow it to fit in the channel. Our observation that the G4 ligand PhenDC_3_ inhibits the transition to the second CMG arrest positions (Figure 3B) favors the latter possibility. Importantly, the channel of the MCM2-7 C-terminal tier contains several protruding loops (Yuan et al., 2016), which could prevent further passage of the G4 and accumulation of the second stalling cluster. Interestingly, similar stalling products (−11 to −24 products) are also observed during DPC repair, only when the DPC is proteolyzed to a peptide (Duxin et al., 2014). This suggests that the “jammed state” might ubiquitously occur, when an impediment fits the N-tier of MCM2-7 but not the C-tier.

### Mechanism of CMG bypass

Our experiments show that CMG can readily bypass a G4 structure on the leading strand template through the action of DHX36. Interestingly, we still observe polymerase stalling at the G4 after CMG bypass, as seen in FANCJ-depletion condition (Figure 4A, lanes 10-15). This raises two possibilities; CMG bypass proceeds without unfolding of the structure, or the G4 structure is unwound but quickly refolds after bypass. We currently favor the first option because biophysical experiments showed that parallel G4 structures require ~200 seconds to fold (Gray et al., 2019). If the G4 structure is not unwound, CMG bypass most likely involves opening of the CMG ring. Such a model is consistent with recent evidence that the MCM2-7 ring opens during replication initiation (Bochman and Schwacha, 2008; Wasserman et al., 2019), and possibly also during DPC repair (Sparks et al., 2019). Strikingly, the requirement of DHX36 or FANCJ for CMG bypass is abolished by a converging fork or when ssDNA is placed downstream of the G4 (Figure 5B, 6D, and S5D). These observations indicate that the primary function of these helicases is to generate ssDNA downstream of the structure, as shown for DPC repair (Sparks et al., 2019), and that the ssDNA triggers CMG bypass via ring-opening by a currently unknown mechanism (Figure S7, iii and iv). Given the directionality and requirements of their ATPase activities (Figure S4A-D), DHX36 likely translocates along the leading strand template, and when it is absent, FANCJ translocates along lagging strand template for ssDNA generation (Figure S7A).

The principle of CMG bypass appears similar between G4 replication and DPC repair, but the mechanism for ssDNA generation past the obstruction differs in important aspects. During DPC repair, CMG bypass requires the 5’-3’ DNA helicase RTEL1 that accumulates at stalled replication forks through PCNA binding (Sparks et al., 2019; Vannier et al., 2013). Unlike DPC repair, DHX36 and FANCJ helicases accumulate at a G4 independently of replication (Figure 4B and S6A). This might be accomplished via their extremely high affinity for G4s (K_D_ of <10 pM for DHX36, ~1 nM for FANCJ) (Giri et al., 2011; Wu and Spies, 2016). In agreement with this idea, a G4 recognition mutant DHX36^Y53A^ does not promote CMG bypass (Figure S4B). While both helicases can create this ssDNA, DHX36 predominates in this process, since depletion of DHX36 enhances the CMG stalling signature while depletion of FANCJ does not (Figure 4A). However, DHX36 is likely inactive before replisome arrival at the G4 structure, as it persists on the G4 in the absence of DNA replication (Figure 4B). We therefore postulate that the replication fork plays a direct role in activating the DHX36 helicase. A backup helicase on the non-G4 strand would be advantageous, for example when DHX36 fails to recognize the structure.

It should be noted that a considerable subset of the G4 structures is bypassed by CMG even when both DHX36 and FANCJ are depleted (Figure 4A) or the G4 is stabilized (Figure 3B), resulting in leading strand extension to the −1 to −3 positions. We therefore assume the existence of yet another bypass pathway. In addition, because G4 structures exist in many different conformations depending on topology and loop size, we speculate that other G4 unwinding mechanisms might exist. Consistently, a recent study suggested that the replication machinery can detect and unwind G4 structure ahead of the replication fork through a Timeless and DDX11 mediated mechanism (Lerner et al., 2020).

### Mechanism of G4 structure unwinding

After CMG bypass, the leading strand polymerase stalls 1 to 3 nucleotides upstream of a G4 structure and is therefore uncoupled from the CMG. Polymerase stalling at the same positions also occurs at the lagging strand G4 after CMG passage. Strikingly, these positions perfectly fit the G4-induced deletion start points found in *C. elegans* (Lemmens et al., 2015), suggesting that polymerase stalling at G4s but not CMG stalling could occasionally lead to DSB formation *in vivo.* Alternatively, this could indicate that G4 structures mostly form on lagging strand templates *in vivo.* While this has been investigated previously, there is no clear direct evidence showing a G4 preference for leading or lagging strand (Lerner and Sale, 2019).

Unlike CMG bypass, DNA synthesis beyond the G4 strictly depends on FANCJ and DHX36, and requires unwinding of the structure (Figure 1D and 4A). Interestingly, CMG bypass is a prerequisite for the efficient FANCJ accumulation and subsequent G4 unwinding, based on the following evidence. First, compromised DNA synthesis past a G4 by defective CMG bypass and FANCJ accumulation in the presence of the ATPase mutant DHX36^E327A^ is rescued by a converging fork, suggesting that CMG bypass is a rate-limiting step for DNA unwinding (Figure 6D and S6A). Second, DNA synthesis past the stabilized G4 takes longer in a single fork than in dual forks, consistent with slower kinetics of CMG bypass in a single fork (Figure 3B). However, it is less clear why CMG bypass is required for FANCJ accumulation. On a lagging strand G4 that does not involve CMG bypass, FANCJ replaces DHX36 with the same kinetics compared to the leading strand (Figure S4F, iii), indicating that CMG bypass per se does not promote the helicase switch. An event that occurs during replication of both strands, but requires CMG bypass on the leading strand, is the collision of the polymerase with the G4 structure (Figure 7). Stalling of the DNA polymerase lowers the affinity for DHX36 that requires a free 3’ end to efficiently bind the G4 while FANCJ does not (Chen et al., 2018; Swan et al., 2009). Therefore, the DNA polymerase could facilitate FANCJ accumulation by preventing re-accumulation of DHX36 that binds to G4s with ~100-fold higher affinity than FANCJ. This model for the helicase switch is consistent with our observation that DNA replication is also required for this switch on a lagging strand G4 (Figure S4F). Future biochemical work is required to establish further details of the helicase switch mechanism.

### G4-induced DNA damage

Our experiments provide direct evidence that a G4 efficiently blocks progression of the vertebrate replisome. G4 formation can be provoked in R-loops at highly transcribed genes (García-Muse and Aguilera, 2016; Hamperl and Cimprich, 2014). Consistently, R-loops were recently shown to drive genomic toxicity of G4 stabilizing ligands (De Magis et al., 2019). G4 ligands also induce synthetic lethality in *BRCA1/2* deficient cancer cells (McLuckie et al., 2013). This strongly indicates that homologous recombination (HR) plays a role in preventing the adverse effects of G4 structures. However, this most likely only comes into play when G4 structures are exceptionally stable which can lead to DSB formation. Under normal circumstances, we assume that the majority of G4 structures can be resolved by an unwinding mechanism such as described here, which proceeds without DSB formation. We thus propose that the DHX36-FANCJ-mediated unwinding represents the primary pathway of G4 maintenance during S phase. Collectively, our work defines a powerful mechanism that maintains genomic stability at G4s. Moreover, specific inhibition of this pathway could be explored for novel therapeutic strategies to treat cancers with deficiencies in DNA repair pathways.

## Acknowledgements

We thank J.C. Walter, M. Tijsterman, and F. Mattirolli for feedback on the manuscript, J.C. Walter and J.L Sparks for pSVRlacO, anti-SPRTN antibody, and methylated M.HpaII, the Hubrecht animal caretakers for animal support, and the other members of the Knipscheer laboratory for discussions. We are grateful to M. Räschle for providing us with the *Xenopus laevis* protein database. K.S. was supported by the Uehara Memorial Foundation, the Mochida Memorial Foundation for Medical and Pharmaceutical Research, and the Japan Society for the Promotion of Science (JSPS) Postdoctoral Fellowship for Research Abroad. N.M.P. was supported by the European Commission and the plasmid pull downs for mass spectrometry, the data analysis, and the production of DHX36 antibodies were funded by the European Union’s Horizon 2020 research and innovation program under the Marie Skłodowska-Curie grant agreement 750035 (ReXeG) to N.M.P. This work was supported by a project grant from the Dutch Cancer Society (KWF HUBR 2015-7736) to P.K. and the Gravitation program CancerGenomiCs.nl from the Netherlands Organisation for Scientific Research (NWO), part of the Oncode Institute, which is partly financed by the Dutch Cancer Society. This research was part of the Netherlands X-omics Initiative and partially funded by NWO, project 184.034.019.

## Author contributions

K.S. and P.K. designed the experiments and prepared the manuscript. K.S. performed experiments with *Xenopus* egg extracts. N.M.P., H.P., and M.A. performed and analyzed the mass spectrometry data. All of the authors read and approved the manuscript.

## Declaration of interests

The authors declare no competing interests.

## METHODS

### Xenopus laevis

Egg extracts were prepared using eggs from adult *Xenopus laevis* female frogs (aged >2 years, Nasco Cat# LM00535) and demembranated sperm chromatin was prepared from the testes of adult *Xenopus laevis* male frogs (purchased from the European Xenopus Resource Centre). All animal procedures and experiments were performed in accordance with national animal welfare laws and were reviewed by the Animal Ethics Committee of the Royal Netherlands Academy of Arts and Sciences (KNAW). All animal experiments were conducted under a project license granted by the Central Committee Animal Experimentation (CCD) of the Dutch government and approved by the Hubrecht Institute Animal Welfare Body (IvD), with project license number AVD80100201711044. Sample sizes were chosen based on previous experience, randomization and blinding are not relevant to this study.

### Insect cell line

Sf9 cells (Thermo Fisher, Cat#B82501) were cultured at 27°C for overexpression of *Xenopus laevis* (xl) FANCJ. Insect cells were cultured in Sf-900 III SFM medium (Thermo Fisher, Cat#12658019).

### Bacterial strains

*E.coli* LOBSTR (DE3) strain (Kerafast, Cat#EC1001) was used for overexpression of *xl*DHX36. *E.coli* BL21 (DE3) strain (New England BioLabs, Cat#C2527) was used for overexpression of xlGeminin and the N-terminal residues (1-170) of xlDHX36. *E.coli* T7 Express strain (New England BioLabs, Cat#C2566) was used for overexpression of LacI and M.HpaII. *E.coli* XL1-Blue strain (Agilent Technologies, Cat#200249) was used for single-stranded (ss) DNA plasmid preparation. *E.coli* cells were cultured in LB medium.

### Preparation of plasmid substrates

Single-stranded DNA (ssDNA) plasmids containing a site-specific G4 motif (5’-GGGAGGGTGGGAGGG-3’) or a control motif (5’-GGGA<u>CCC</u>TGGGAGGG-3’) were prepared by viral replication using a helper phage M13K07 (New England BioLabs, Cat#N0315S). *E.coli* XL1-Blue cells were transformed with pBluescript SK(-) plasmid containing the motif and cultured in 50 mL LB medium supplemented with ampicillin (100 μg/mL final concentration) at 37°C until OD_600_ reaches ~0.05. 50 μL M13K07 (1×10^11^ pfu/mL) phage was then added, and the cells were cultured for further 18 hours in the presence of kanamycin (70 μg/mL final concentration). The medium was collected after centrifugation (4,000 g) for 10 minutes and mixed with 0.2 volumes of PEG solution (2.5 M NaCl and 20% PEG-8000) for an hour at 4°C. After centrifugation (12,000 g) for 10 minutes at 23°C, the pellet was resuspended in 1.6 μL TE (10 mM Tris-HCl pH 8 and 1 mM EDTA) and centrifuged by 17,000 g for a minute at 23°C to remove any remaining cells. The supernatant was collected, after which 0.2 volumes of PEG solution was added and the solution was centrifuged by 17,000 g for 10 minutes at 23°C. The pellet was resuspended with 300 μL TE and purified by phenol/chloroform extraction. Purified DNA was then ethanol-precipitated with 20 μg glycogen, resuspended with 25 μL TE, and stored at −80°C.

The double-stranded (ds) plasmids containing a DNA interstrand crosslink derived from cisplatin (pICL^Pt^) was prepared as described (Enoiu et al., 2012). dsDNA plasmids containing a G4 motif were prepared by a similar method. First, short DNA duplexes were generated by heating a pair of oligonucleotides (Extended Data Table 1) to 80°C for 5 minutes and slowly cooling down to room temperature in ~2 hours in annealing buffer (10 mM Tris-HCl pH and 25 mM KCl). To generate pdsG4^BOT^ and pdsCON, oligonucleotides *A* and *B,* or *C* and *D* was used, respectively. For pPolyT, pCON, pG4^BOT^, pG4^DOWN^, and pG4^UP^, oligonucleotides *E* and either *F, G, H, I,* or *J* were used, respectively. For pG4^TOP^, oligonucleotides *K* and *L* were used. The resulting duplexes were ligated into the BbsI sites of the pSVR*lacO* vector (Zhang et al., 2015). After ligation, the closed circular plasmid was purified using a cesium chloride gradient ultracentrifugation, followed by butanol extraction, concentration and buffer exchange to TE with an Amicon Ultra-4 centrifuge filter unit (30 kDa cut-off, Merck Millipore, Cat#UFC803024).

To make a pG4^Lead-DPC^, pKS was created by replacing the BbsI fragment of pSVR*lacO* with a duplex, consisting of oligonucleotides *M* and *N,* containing two Nt.BbvCI nicking sites. pKS was nicked with Nt.BbvCI and purified using Wizard SV Gel and PCR Clean-Up System (Promega, Cat#A9281). The resulting short ssDNA fragment was then replaced with a 5-fluoro-2’-deoxycytidine (FdC)-modified oligonucleotide (400 nM) by heating the mixture (containing 50 nM nicked pKS) to 80°C for 5 minutes, and cooled down to room temperature in ~2 hours. To avoid re-annealing of the original fragment, an excess (2.4 μM) of oligonucleotide *O* (complementary to the original fragment) was added. The annealed fragment was ligated with T4 DNA ligase (66.7 unit/μL at a final concentration, New England BioLabs, Cat#M0202M) by incubating in reaction buffer (50 mM Tris-HCl pH7.5, 10 mM MgCl2, 4 mM ATP, and 10 mM dithiothreitol) for 18 hours at 16°C. The modified plasmid was purified and concentrated to ~150 ng/μL with Wizard SV Gel and PCR Clean-Up System using elution buffer (10 mM Tris-HCl pH 7.5, 10 mM KCl, and 0.1 mM EDTA). To induce a covalent crosslink between FdC and the catalytic cysteine of DNA methyltransferase M.HpaII (Extended Data Fig. 6c) (Chen et al., 1991), the purified plasmid was mixed with His6-tagged M.HpaII, that was methylated on lysine residues to prevent proteolysis in *Xenopus* egg extract (a kind gift from J.C. Walter) (Larsen et al., 2019), for 18 hours at 37°C in crosslink buffer (50 mM potassium acetate, 20 mM Tris-acetate, 10 mM Magnesium acetate, 0.1 mg/mL bovine serum albumin (BSA), 100 μM S-adenosylmethionine, 30 mM KCl, 9% glycerol, and 0.3 mM dithiothreitol, pH 7.9).

### *Xenopus* egg Extracts and DNA Replication

*Xenopus laevis* female frogs were used as a source of eggs. Egg production, preparation of *Xenopus* high-speed supernatant (HSS), demembranated sperm chromatin, and nucleoplasmic extract (NPE), and DNA replication were performed as previously described (Sparks and Walter, 2019; Walter et al., 1998).

For replication of ssDNA templates, ssDNA plasmids (20 ng/μL final concentration) were incubated with primer A (1.0 μM final concentration) in annealing buffer for 5 minutes at 80°C and cooled down to room temperature in ~2 hours to allow the G4 structure to form and the primer to anneal. Where indicated, the G4 stabilizing compound PhenDC_3_ was added to the mixture at 50°C and incubated for 30 minutes after which the mixture was further cooled down to room temperature. To prevent 5’ to 3’ DNA resection in egg extract, the primer was synthesized with the twelve most 5’ nucleotides connected by phosphorothioate bonds. To initiate replication, the primed template (2.7 ng/μL final concentration) was added to HSS supplemented with 3 ng/μL nocodazole, 18.9 mM phosphocreatine, 1.9 mM ATP, and 4.7 ng/μL creatine phosphokinase at room temperature. For nascent strand labelling, HSS was supplemented with ^32^P-a-dCTP. For extraction of replicated samples, aliquots of the reaction (5 μL) were stopped with 45 μL stop solution II (50 mM Tris pH 7.5, 0.5% SDS, and 10 mM EDTA pH 8.0) at the indicated time points. Samples were then treated with RNase (0.15 mg/mL) for 30 minutes at 37°C, followed by Proteinase K (0.5 mg/mL) treatment overnight at room temperature. DNA was phenol/chloroform extracted, ethanol precipitated with glycogen (20 μg), and resuspended in 5 μL TE.

For replication of dsDNA templates, the G4 structure was induced on dsDNA plasmids (75 ng/μL final concentration) prior to replication by the same method as for the ssDNA template. For replication of pG4^BOT-CON^ and pG4^Lead-DPC^, replication was conducted in a SPRTN depleted background to prevent M.HpaII degradation in *Xenopus* egg extracts (Larsen et al., 2019). The plasmids (15 ng/μL for plasmid pull-down assay and 9 ng/μL for all other assays) were then incubated with HSS for 20 minutes at room temperature to assemble pre-replication complexes. Two volumes of NPE (diluted to 40% with ELBS buffer containing 10 mM HEPES-KOH pH 7.7, 50 mM KCl, 2.5 mM MgCl2, and 250 mM sucrose) were then added to fire a single round of DNA replication. For nascent strand labelling, NPE was supplemented with ^32^P-α-dCTP. To inhibit replication initiation, recombinant *xl*Geminin (400 nM final concentration) was added to HSS and incubated for 10 minutes at room temperature prior to DNA addition. To block CMG unloading, an inhibitor of the p97 segregase, NMS-873 (Sigma-Aldrich, Cat#SML1128-5MG, 200 μM final concentration), was added to NPE and incubated for 10 minutes at room temperature prior to mixing with HSS. For replication in the presence of LacI, plasmids were incubated with 1.33 volumes of 30 μM biotinylated LacI for 30 minutes at room temperature prior to HSS addition.

Where indicated, LacI was released from DNA by adding isopropyl β-D-1-thiogalactopyranoside (IPTG, 9 mM final concentration) to replication reaction. The 0 min time point was taken immediately after NPE addition. DNA was extracted from replication reactions by the same method as described for the ssDNA templates. To monitor the replication fork blockage by LacI bound to the *lacO* locus, extracted replication products were digested with HincII for 3 hours at 37°C and separated on a 0.8% agarose gel in 1x TBE buffer (89 mM Tris-borate and 2 mM EDTA). DNA was visualised by autoradiography using Typhoon TRIO+ (GE Healthcare).

### Plasmid pull-down mass spectrometry

Replicating ssDNA plasmids were pulled down as previously described (Budzowska et al., 2015; Larsen et al., 2019). Streptavidin magnetic beads (Dynabeads M-280, Thermo Fisher, Cat#DB11205) were washed 3 times with 1 volume of wash buffer I (50 mM Tris-HCl pH7.5, 150 mM NaCl, 1 mM EDTA, and 0.02% Tween-20), resuspended with 1 volume of wash buffer I, and incubated with biotinylated LacI (2 μM final concentration) for 45 minutes with mixing every 10 minutes at room temperature. The beads were washed 4 times with 1 volume of IP buffer (ELBS buffer supplemented with 0.25 mg/mL BSA and 0.02% Tween 20), resuspended with 6.64 volume of IP buffer, and stored at 4°C. To capture replication intermediates via LacI that non-specifically binds to ssDNA and dsDNA (Riggs et al., 1972), 150 μL of the replication reaction was mixed with 750 μL beads solution (containing 113 μL biotin-LacI-bound Streptavidin magnetic beads) at the indicated times, and incubated for 30 minutes at 0-2°C with mixing every 10 minutes. The beads were washed three times with 750 μL IP buffer containing 0.03% Tween 20, and resuspended in 40 μL 1x SDS sample buffer (75 mM Tris-HCl pH6.8, 10% glycerol, 2.5% SDS, 10% (v/v) Bond-Breaker TCEP Solution (Thermo Fisher, Cat# 7772), and 0.02% (w/v) bromophenol blue). The samples were heated at 95°C for 5 min and separated on a 12% Bis-Tris SDS-PAGE gel (Biorad).

### Mass spectrometry

#### Data collection

The gel was run for 2-3 cm and stained with colloidal coomassie dye G-250 (Gel Code Blue Stain Reagent, Thermo Scientific) after which each lane was cut out. Gel pieces were reduced, alkylated and digested overnight with trypsin at 37°C. The peptides were extracted with 100% acetonitrile (ACN) and dried in a vacuum concentrator. Samples were resuspended in 10% (v/v) formic acid for UHPLC-MS/MS. The data was acquired using an UHPLC 1290 system coupled to an Orbitrap Q Exactive Biopharma HF mass spectrometer (Thermo Scientific). Samples were first trapped (Dr Maisch Reprosil C18, 3 μm, 2 cm x 100 μm) before being separated on an analytical column (Agilent Poroshell EC-C18, 278 μm, 40 cm x 75 μm), using a gradient of 100 min at a column flow of 300 nl/min. Trapping was performed at 5 μL/min for 5 min in solvent A (0.1 % formic acid in water) and the gradient was as follows 13-44% solvent B (0.1% formic acid in 80% acetonitrile) in 95 min, 44-100% in 3 min, 100% solvent B for 1 min and 100-0% in 1 min. Full scan MS spectra from m/z 375 – 1600 were acquired at a resolution of 60,000 at m/z 400 after accumulation to a target value of 3e6. Up to ten most intense precursor ions were selected for fragmentation. HCD fragmentation was performed at normalised collision energy of 27% after the accumulation to a target value of 1e5. MS/MS was acquired at a resolution of 30.000.

#### Data analysis

Raw data were analyzed with the MaxQuant software (version 1.5.0.17) using label-free quantification (Cox and Mann, 2008). A false discovery rate (FDR) of 0.01 and a minimum peptide length of 7 amino acids were used. MS/MS spectra were searched against a non-redundant *Xenopus* database (Temu et al., 2016). For the Andromeda search the enzyme trypsin was chosen allowing for cleavage N-terminal to proline. Cysteine carbamidomethylation was selected as a fixed modification, and protein N-terminal acetylation and methionine oxidation were selected as variable modifications. Two missed cleavages were allowed maximally. Initial mass deviation of precursor ion was up to 7 ppm and mass deviation for fragment ions was 0.05 Dalton. Protein identification required one unique peptide to the protein group and “Match between run” was enabled.

#### Statistical analysis

All bioinformatics analysis was carried out with the Perseus software Version 1.6.10.0. For each comparison, the processed data was filtered to contain at least 3 valid values in at least one of the replicate group (four repeats per condition).

### Antibodies and immunodepletion

Antibodies against *xl*FANCJ (Castillo Bosch et al., 2014)*, xl*MCM7 (Walter and Newport, 2000), *xl*PCNA (Kochaniak et al., 2009), *xl*SPRTN (Larsen et al., 2019), and histone H3 (Abcam, Cat#ab1791) were previously described. The *xl*DHX36 antibody was raised against N-terminal residues (1-170) of *xl*DHX36. The cDNA encoding the fragment was codon-optimized for *E. coli,* synthesized (gBlocks Gene Fragments, Integrated DNA Technologies), and ligated into the BamHI-XhoI sites of the pETDuet-1 vector (Novagen, Cat#71146-3). The fragment was overexpressed in *E. coli* BL21(DE3) cells with a N-terminal His6-tag, and purified by the method described previously (Brandsma et al., 2019). The purified antigen was used for immunization of rabbits (PRF&L, Canadensis, USA). Specificity of the antisera was confirmed using immunoblotting. Antibodies used for ChIP experiments were purified with rProtein A Sepharose (PAS) beads (GE Healthcare, Cat#171279-01).

For depletion, one volume of Dynabeads™ Protein A beads (Thermo Fisher Scientific, Cat#10008D) was pre-incubated with half volumes of antibody for 30 minutes at room temperature. For mock depletion, pre-immunized rabbit serum was used. To deplete HSS and NPE, one volume of each antibody-bound beads was incubated with one-half volumes of extract for 30 minutes at room temperature for two rounds.

### Protein purification

Recombinant C-terminal FLAG-tagged *xl*FANCJ (Castillo Bosch et al., 2014), *xl*Geminin (McGarry and Kirschner, 1998), biotinylated LacI (Dewar et al., 2015), and methylated M.HpaII (Larsen et al., 2019) were prepared as previously described. The *xl*FANCJ^K52R^ mutant was created by a site-directed mutagenesis using oligonucleotides *K52R for* and *K52R rev* (Extended Data Table 1) and purified by the same method as wild-type *xl*FANCJ.

For purification of *xl*DHX36, the cDNA encoding full-length *xl*DHX36 (gBlocks Gene Fragments, Integrated DNA Technologies) were ligated into the BamHI-XhoI sites of the pETDuet-1 vector, and the protein were overexpressed as a N-terminal His6-tagged protein in the *E.coli* LOBSTR (DE3) cells cultured in 4 L LB medium, as previously described (Takahashi et al., 2014). The cells were collected, resuspended in buffer A (50 mM Tris-HCl pH 8.0, 10% glycerol, 0.5 M NaCl, 1 mM phenylmethylsulfonyl fluoride, 20 mM imidazole, 0.1% Tween-20 and 5 mM dithiothreitol), and disrupted by sonication. The supernatant was then separated from the cell debris by centrifugation (39,191 g) for 25 minutes at 4°C, and treated with polyethyleneimine (0.05%, v/v). After centrifugation (16,639 g) for 10 minutes at 4°C, the supernatant was mixed with nickel-nitrilotriacetic acid (Ni-NTA) agarose resin (1.2 mL, Thermo Fisher, Cat#R90115) at 4°C for an hour. The Ni-NTA beads were packed into a disposable chromatography column (Bio-Rad, Cat#731-1550), and were washed with 60 mL buffer A, followed by 30 mL buffer B (50 mM Tris-HCl pH 8.0, 10% glycerol, 250 mM NaCl, 20 mM imidazole, and 5 mM dithiothreitol). His6-tagged *xl*DHX36 was eluted with a 8 mL buffer B containing 400 mM imidazole, and loaded on 1 mL HiTrap Heparin HP (GE Healthcare, Cat# 17040601) equilibrated with buffer C (20 mM Tris-HCl pH 8.0, 10% glycerol, 250 mM NaCl, and 5 mM dithiothreitol). The column was subsequently washed with buffer C, and the protein was eluted with a 60 mL linear gradient of 250 to 1000 mM NaCl in buffer C. Peak fractions were collected, concentrated with Amicon Ultra-4 Centrifugal Filter Unit (100 kDa cut-off, Merck Millipore, Cat#UFC810024) to ~1 mg/mL, snap-frozen, and stored at −80°C. The *xl*DHX36 mutants were created by a site-directed mutagenesis using oligonucleotides *Y53A for* and *Y53A rev* (for DHX36^Y53A^) or *E327A for* and *E327A rev* (for DHX36^E327A^) and purified by the same method as the wild-type protein. For rescue experiments, depleted NPE (for dsDNA template replication) or HSS (for ssDNA template replication) was supplemented with 35 nM recombinant *xl*FANCJ or 250 nM recombinant *xl*DHX36. The concentration of the recombinant proteins was determined by SDS-PAGE with Coomassie Brilliant Blue staining, using BSA as a standard protein.

### Nascent strand analysis

Nascent strands on ssDNA templates were analysed as previously described (Castillo Bosch et al., 2014). Extracted replication products were mixed with the same volume of Gel Loading Buffer II (Thermo Fisher, Cat#AM8547), heated for 4 minutes at 98°C, snap-cooled on ice, and separated on 6% urea-PAGE gels, which were subsequently dried and exposed to a phosphor screen. DNA was visualized using a Typhoon TRIO+.

Nascent strands on dsDNA templates were analysed as previously described (Sato et al., 2020). Extracted replication products were digested with HincII, or HincII and ClaI, for 3 hours at 37°C, ethanol-precipitated, and resuspended in 12 μL alkaline loading buffer (50 mM NaOH, 2.5 % Ficoll-400, and 1 mM EDTA). Fragments were then separated on a 0.8% agarose gel in alkaline buffer (50 mM NaOH and 1 mM EDTA), after which the gel was dried on Amersham Hybond-XL membrane (GE Healthcare, Cat# RPN203S) and exposed to a phosphor screen.

To determine exact replication stalling positions, extracted replication products were analysed by sequencing gel as previously described (Räschle et al., 2008). The samples were digested either with AflIII and BamHI or with AflIII, BamHI and ClaI for 3 hours at 37°C, mixed with the same volume of Gel Loading Buffer II, heated at 98°C for 5 minutes, snap-cooled on ice for 5 minutes, and separated on a 7% urea-polyacrylamide sequencing gel prepared in 0.8x TTE buffer (71 mM Tris, 23 mM taurine, and 0.4 mM EDTA, pH 8.9). After gel drying, the products were visualized by autoradiography. The size of the stalling products was determined using sequencing ladders and PCR-amplified “-1” and “-2” fragments. The ladders were generated using pdsG4^BOT^ and *primer T* by the Thermo Sequenase Cycle Sequencing kit (Thermo Fisher, Cat#785001KT) according to the manufacturer’s protocol. −1 and −2 products on bottom strand were prepared by PCR using pdsG4^BOT^ and *primer T* labelled with ^32^P at the 5’-end and either *primer T-1* or *primer T-2,* respectively.

### Plasmid pull-down assay

Replicating dsDNA plasmids were pulled down as previously described (Budzowska et al., 2015). At the indicated times, 10 μL replication samples were mixed with 7.5 μL biotin-LacI-bound Streptavidin magnetic beads suspended in 50 μL IP buffer containing 0.03% Tween 20, and incubated for 30 minutes at 0-2°C without pipeting. The beads were washed three times with 75 μL IP buffer, and suspended in 20 μL 1x SDS sample buffer. Plasmid-bound proteins were then separated by SDS-PAGE and visualized by Western blot using the indicated antibodies.

### Two-dimensional gel electrophoresis (2DGE)

2DGE was performed as previously described (Long et al., 2011). Extracted replication samples were digested with HincII for 3 hours at 37°C, and analyzed by two consecutive electrophoreses. For the first dimension, the HincII digested samples were separated with 0.4% agarose gel in 0.5x TBE buffer at 0.86 V/cm for 24 hours at room temperature. The lanes of interest were cut out, casted across the top of the second-dimension gel consisting of 1% agarose with 0.3 μg/mL ethidium bromide, and run in 0.5x TBE buffer containing 0.3 μg/mL ethidium bromide with buffer circulation at 3.5 V/cm for 14.5 hours at room temperature. The gel was dried on Amersham Hybond-XL membrane and exposed to a phosphor screen. DNA was visualized using a Typhoon TRIO+.

### Chromatin immunoprecipitation (ChIP)

ChIP was performed similar to described previously (Pacek et al., 2006). At the indicated times, replication samples (3 μL) were crosslinked with 47 μL ELBS buffer containing 1% formaldehyde for 10 minutes at room temperature. An unrelated non-damaged control plasmid (pQuant, 0.5 ng/μL) was co-incubated with HSS to be used as an internal control for quantifications. After quenching the formaldehyde by addition of 5 μL 1.25 M glycine, the samples were passed through a Micro Bio-Spin 6 Chromatography column (Bio-Rad, Cat#7326222), sonicated, and immunoprecipitated with the indicated antibodies (5 μg) bound to PAS beads. The protein-bound DNA fragments were eluted with ChIP elution buffer (50 mM Tris pH7.5, 10 mM EDTA, and 1% SDS) and the crosslinks were reversed by consecutive incubation for 6 hours at 42°C and then for 9 hours at 70°C. DNA was phenol/chloroform-extracted, followed by quantitative PCR in 10 μl reaction buffer (6 mM Tris pH 8.3, 25 mM KCl, 2.5 mM MgCl2, 0.3 mM dNTPs, 0.1% Tween-20, 0.1 mg/ml BSA, 1:66,500 SYBR Green I (Sigma-Aldrich, Cat#S9430), and Hot Start Taq DNA polymerase), using 0.25 μM of following primer pairs (Extended Data Table 1): *G4 for* and *G4 rev* (for G4 locus, 37-136 bp upstream from G4s), *lacO for* and *lacO rev* (for *lacO* locus, 295-388 bp downstream from Gs), and *pQuant for* and *pQuant rev* (for assessment of background binding of the proteins on pQuant). The values from *pQuant* primers were subtracted from the values for *G4* and *lacO* primers.

## Notes

### Competing Interest Statement

The authors have declared no competing interest.

