## Supplemental Figures for "Multistep mechanism of DNA replication-coupled G-quadruplex resolution"

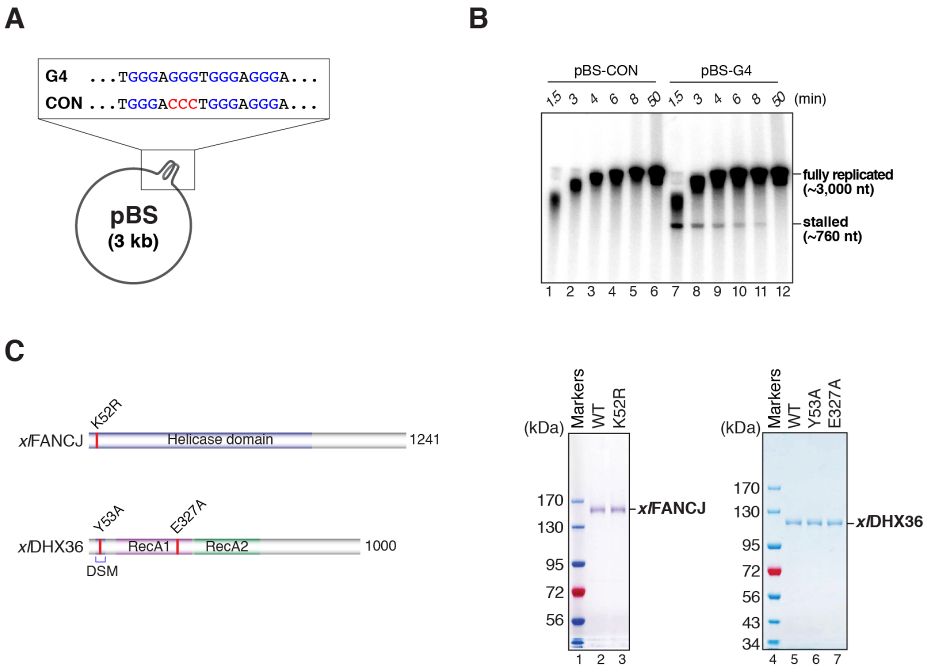


**Figure S1. Replication of ssDNA plasmids in *Xenopus* egg extract, Related to Figure 1**

(A) Schematic representation of pBS-G4 and pBS-CON showing the G4 and control sequences. G- and C-tracts are colored in blue and red, respectively.

(B) pBS-CON and pBS-G4 were replicated in HSS in the presence of $^{32}$P-$\alpha$-dCTP (see also Figure 1A). Replication products were extracted, separated by denaturing PAGE, and visualized by autoradiography. Nascent strands stalled at the G4 sequence ('stalled') and fully replicated strands ('fully replicated') are indicated.

(C) Schematic representation of the *Xenopus laevis* (*xl*) FANCJ and DHX36 proteins. The helicase domain of FANCJ, the DHX36-specific motif (DSM) that recognizes G4 structures, and two RecA-like domains (RecA1 and RecA2) that mediate DHX36's ATPase activity are indicated. The residues mutated in FANCJ$^{K52R}$, DHX36$^{Y53A}$, and DHX36$^{E327A}$ are indicated with red bars.

(D) Purified wild-type (WT) FANCJ, FANCJ$^{K52R}$, WT DHX36, DHX36$^{Y53A}$, and DHX36$^{E327A}$ used in this study, were analyzed by PAGE with Coomassie Brilliant Blue staining alongside molecular-weight markers.

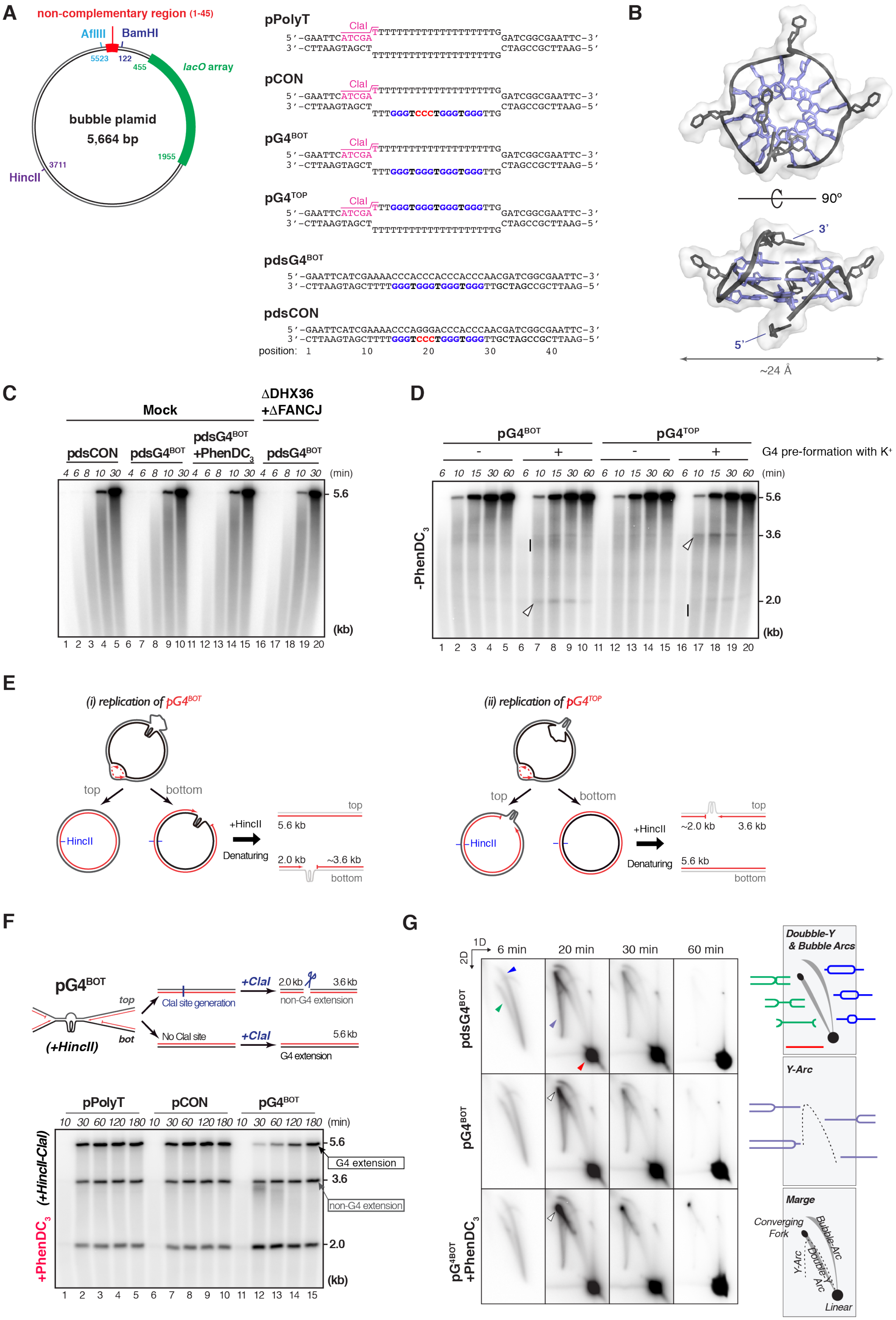
A
non-complementary region (1-45)
AflIII
BamHI
5523
122
455
lacO array
bubble plamid
5,664 bp
1955
3711
HincII
pPolyT
ClaI
5'-GAATTCATCGATTTTTTTTTTTTTTTTTTTTGATCGGCGAATTC-3'
3'-CTTAAGTAGCTTTTTTTTTTTTTTTTTTTTTCTAGCCGCTTAAG-5'
pCON
5'-GAATTCATCGATTTTTTTTTTTTTTTTTTTTGATCGGCGAATTC-3'
3'-CTTAAGTAGCTTTTGGGTCCCTGGGTGGGTTGCTAGCCGCTTAAG-5'
pG4BOT
5'-GAATTCATCGATTTTTTTTTTTTTTTTTTTTGATCGGCGAATTC-3'
3'-CTTAAGTAGCTTTTGGGTGGGTGGGTGGGTTGCTAGCCGCTTAAG-5'
pG4TOP
5'-GAATTCATCGATTGGGTGGGTGGGTGGGTTGGATCGGCGAATTC-3'
3'-CTTAAGTAGCTTTTTTTTTTTTTTTTTTTTTCTAGCCGCTTAAG-5'
pdsG4BOT
5'-GAATTCATCGAAAACCCACCCACCCACCCAACGATCGGCGAATTC-3'
3'-CTTAAGTAGCTTTTGGGTGGGTGGGTGGGTTGCTAGCCGCTTAAG-5'
pdsCON
5'-GAATTCATCGAAAACCCAGGGACCCACCCAACGATCGGCGAATTC-3'
3'-CTTAAGTAGCTTTTGGGTCCCTGGGTGGGTTGCTAGCCGCTTAAG-5'
position: 1 10 20 30 40
B
90°
3'
5'
~24 Å
C
Mock
ΔDHX36 +ΔFANCJ
pdsCON
pdsG4BOT
pdsG4BOT +PhenDC3
pdsG4BOT
4 6 8 10 30 4 6 8 10 30 4 6 8 10 30 4 6 8 10 30 (min)
5.6
(kb)
1 2 3 4 5 6 7 8 9 10 11 12 13 14 15 16 17 18 19 20
D
pG4BOT
pG4TOP
- + - +
G4 pre-formation with K+
6 10 15 30 60 6 10 15 30 60 6 10 15 30 60 6 10 15 30 60 (min)
-PhenDC3
5.6
3.6
2.0
(kb)
1 2 3 4 5 6 7 8 9 10 11 12 13 14 15 16 17 18 19 20
E
(i) replication of pG4BOT
top
bottom
HincII
+HincII
Denaturing
top
5.6 kb
2.0 kb
~3.6 kb
bottom
(ii) replication of pG4TOP
top
bottom
HincII
+HincII
Denaturing
top
~2.0 kb
3.6 kb
5.6 kb
bottom
F
pG4BOT
top
bot
(+HincII)
ClaI site generation
+ClaI
2.0 kb
3.6 kb
non-G4 extension
No ClaI site
+ClaI
5.6 kb
G4 extension
pPolyT
pCON
pG4BOT
10 30 60 120 180 10 30 60 120 180 10 30 60 120 180 (min)
(+HincII-ClaI)
+PhenDC3
5.6
G4 extension
3.6
non-G4 extension
2.0
(kb)
1 2 3 4 5 6 7 8 9 10 11 12 13 14 15
G
1D
2D
6 min
20 min
30 min
60 min
pdsG4BOT
pG4BOT
pG4BOT +PhenDC3
Doubble-Y & Bubble Arcs
Y-Arc
Marge
Converging Fork
Bubble-Arc
Double-Y Arc
Y-Arc
Linear

**Figure S2. Replication fork stalling at G4 structures, Related to Figure 2**

(A) Schematic representation of dsDNA plasmids used in this study. Inserted (non)complementary duplex sequences containing a G4 motif or control motif are indicated (right panel). A unique ClaI restriction site (magenta), G-tracts (blue), and C-tracts (red) are indicated. Restriction sites used in this study and the site of a 48-repeat *lacO* array are indicated with their positions on the plasmid (left panel).

(B) NMR solution structure of the G4, of which the sequence was used for the plasmids described in (A) (PDBID:2LK7) (Do and Phan, 2012). Guanine and thymine bases are colored as in (A). Bottom panel: The structure rotated by 90º. Diameter is indicated.

(C) pdsCON and pdsG4$^{BOT}$ were replicated in mock- or DHX36-FANCJ-depleted extracts in the presence of $^{32}$P-$\alpha$-dCTP. Replication products were isolated, linearized with HincII which cuts 2.0 kbp upstream from the G4, and analyzed by denaturing agarose gel electrophoresis and autoradiography. Where indicated, pdsG4$^{BOT}$ was treated with PhenDC$_3$ prior to replication. Replication arrest at the G4 is expected to yield digestion products of 2.0 kb and 3.6 kb. However, full-length replication products (5.6 kb) readily accumulated and no intermediates were observed, indicating that the G4 structure does not efficiently form in any of the conditions.

(D) pG4$^{BOT}$ and pG4$^{TOP}$ were replicated in the presence of $^{32}$P-$\alpha$-dCTP and analyzed as in (C). Where indicated, G4 formation was suppressed by adding lithium ion instead of potassium ion. 2.0 kb and ~3.6 kb nascent products observed during pG4$^{BOT}$ replication are indicated with an arrowhead and a bar, respectively. 3.6 kb and ~2.0 kb nascent products observed during pG4$^{TOP}$ replication are indicated with a bar and an arrowhead, respectively. While replication of plasmids in the presence of potassium showed transient accumulation of ~2.0 kb and ~3.6 kb digestion products (lanes 6-10 and lanes 16-20), replication of the plasmids in the absence of potassium immediately yielded the full-length 5.6 kb product without accumulation of intermediates (lanes 1-5 and lanes 11-15). These data indicate that replication stalling occurs at the G4 structure.

(E) Model for replication of pG4$^{BOT}$ and pG4$^{TOP}$. (i) For pG4$^{BOT}$, upon replication stalling at the G4 structure, a 2.0 kb nascent product containing the 3' end and a ~3.6 kb nascent product containing the 5' end are formed on the bottom strand template after digestion with HincII. Since the free 5' end is prone to degradation in the extract, the latter product is less defined in size. The top strand template is readily replicated without stalling, generating a 5.6 kb nascent product after the digestion. (ii) Analogous products are generated during pG4$^{TOP}$ replication, in which stalled nascent strands are detected as discrete 3.6 kb and less defined ~2.0 kb fragments.

(F) pPolyT, pCON, and pG4$^{BOT}$ were replicated in the presence of $^{32}$P-$\alpha$-dCTP and PhenDC$_3$. Replication products were isolated and digested with HincII and ClaI and analyzed as in (C). Digested nascent extension products of pG4$^{BOT}$ are depicted (Schematic, upper panel). Since a ClaI restriction site is designed in the ssDNA-dsDNA junction, ClaI digests the top strand only after the motif is replicated (Figure S2A). Thus, the extension product of the non-G4 template strand is detected as 3.6-kbp (gray open box) and 2.0-kbp fragments in pG4$^{BOT}$ replication, while the extension product of the G4 template strand is detected as a 5.6-kbp product (black open box). During replication of pG4$^{BOT}$, the non-G4 template strand is replicated with comparable kinetics to control plasmids, whereas replication of the G4 template strand is specifically compromised.

(G) pCON and pG4$^{BOT}$ were replicated in the presence of $^{32}$P-$\alpha$-dCTP with or without PhenDC$_3$. Replication products were isolated, linearized with HincII, and analyzed by two-dimensional gel electrophoresis (2DGE). The right cartoon illustrates 2DGE patterns of relevant DNA intermediates. Replicated pCON migrated mostly as a linear product after 30 minutes (red arrowhead). Although replicated pG4$^{BOT}$ moderately persisted at an apex of the double Y-arc at 20 minutes (white arrowheads), most intermediates migrated as linear products by 30 minutes, even in the presence of PhenDC$_3$, when replication stalling was prominent (Figure 2B). These data suggest that replication forks stall at the G4 structure, but the intermediate is quickly resolved into daughter molecules.

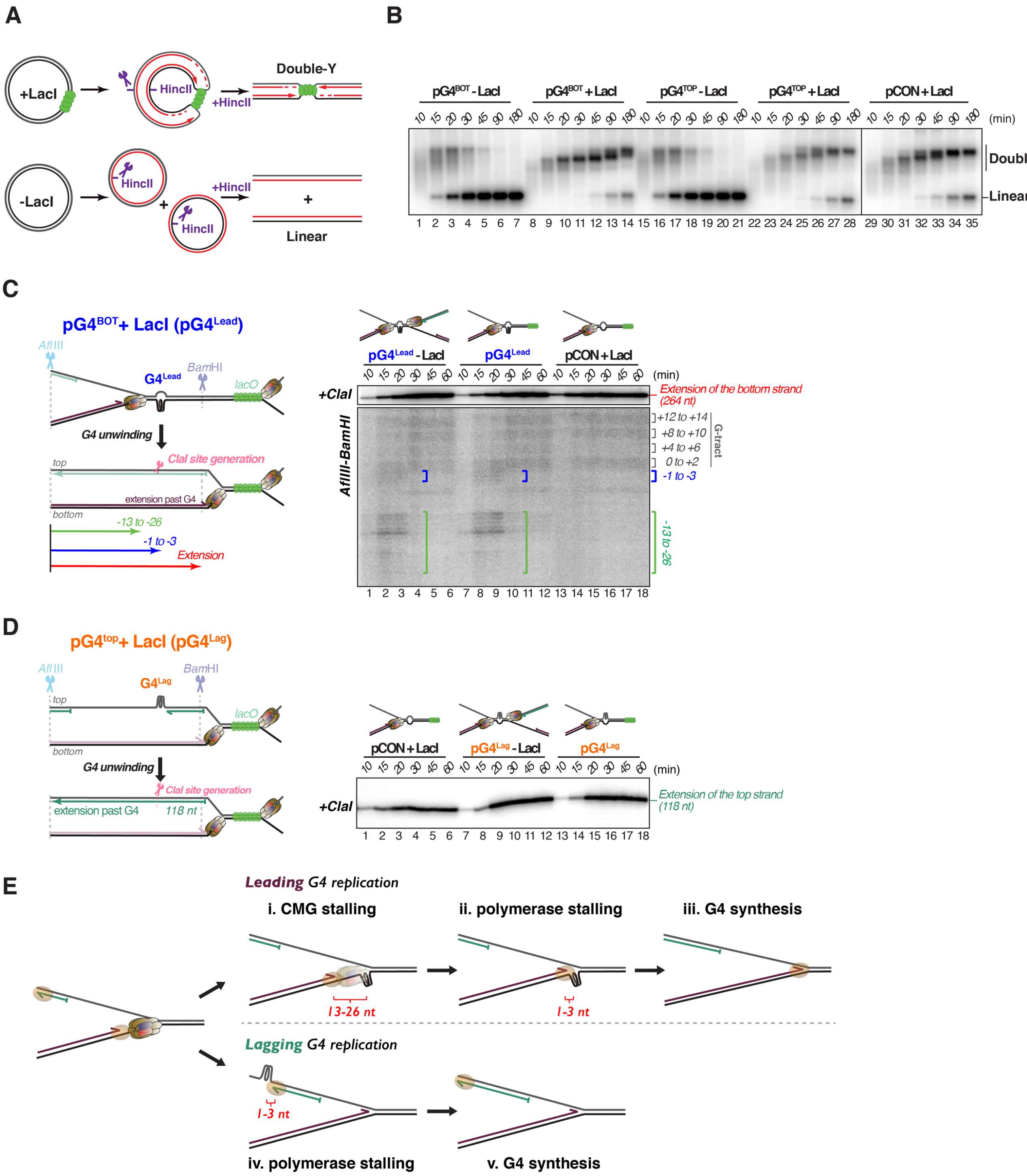


**Figure S3. Replication fork convergence is not required for G4 resolution, Related to Figure 3**

(A) Schematic representation of replication intermediates of plasmids containing a *lacO* array in the presence and absence of LacI (green spheres) after digestion with HincII.

(B) pG4$^{BOT}$, pG4$^{TOP}$, and pCON were replicated in the presence of $^{32}$P-$\alpha$-dCTP with or without LacI. Replication products were isolated, digested with HincII, separated on an agarose gel, and visualized by autoradiography. Replication forks were efficiently blocked and only slowly released after one hour indicated by the appearance of linear molecules.

(C) Schematic representation of nascent leading and lagging strand extension products generated when $pG4^{Lead}$ is replicated (left panel). AflIII and BamHI digestion sites are indicated. Digestion with ClaI specifically digests the extension products of the top strand. In the presence of LacI, extension products of the top strand correspond to lagging strands, and extension products of the bottom strand correspond to leading strands. $pG4^{Lead}$ and LacI-bound pCON were replicated in the presence of $^{32}P$-$\alpha$-dCTP. Where indicated, LacI was omitted. Replication products were digested with AflIII, BamHI and ClaI (upper gel) or AflIII and BamHI (lower gel), separated on a denaturing polyacrylamide gel, and visualized by autoradiography (right panel). Nascent leading strand products are indicated with brackets with their stalling positions printed next to them. The -1 product corresponds to a product in which synthesis has stopped 1 nt before the first guanine of the G4 structure. Two stalling clusters (blue and green brackets) were specifically generated during replication of the G4 structure. Replication also stalled at G-tracts within the G4 motif (black brackets), but this is not caused by the G4 structure as similar replication products were generated during replication of LacI-bound pCON (see also Figure 3B). Brown hexamer, CMG; Green sphere, LacI.

(D) Schematic representation of nascent leading and lagging strand extension products generated when $pG4^{Lag}$ is replicated (left panel). $pG4^{Lag}$ and LacI-bound pCON were replicated in the presence of $^{32}P$-$\alpha$-dCTP. Where indicated, LacI was omitted. Replication products were digested with AflIII, BamHI and ClaI, and analyzed as in (C) (right panel). Accumulation of the 118 nt product indicates the extension past the G4 structure by lagging strand synthesis.

(E) Model of step-wise replication of a G4 structure on the leading and lagging strand templates. Upon encounter of a G4 structure, leading strand synthesis transiently stalls at 13 to 26 nt upstream of the structure, and comes to second arrest at 1 to 3 nt upstream of the structure (top). Given the stalling positions from the G4 structures and the transient MCM7 accumulation at the structure (Figure 3D), we conclude that the first stalling products result from CMG stalling at the G4 structure (i) and the second from DNA polymerase stalling at the G4 (ii). The G4 structure is subsequently unwound and replicated (iii). In contrast, the lagging strand directly stalls at 1 to 3 nt upstream of the G4 structure (iv), followed by G4 resolution and synthesis (v, bottom).

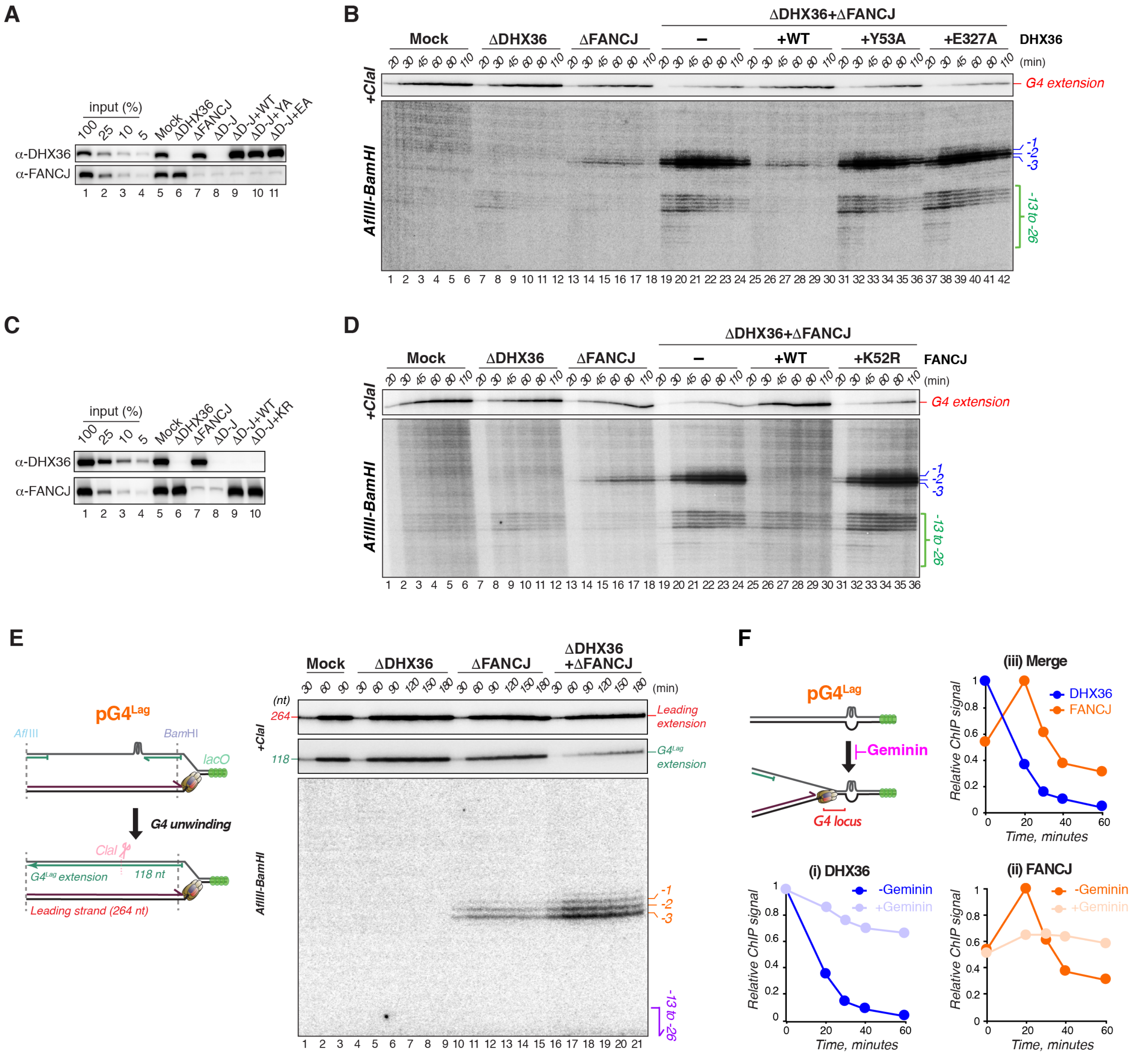


**Figure S4. DHX36 and FANCJ are essential for efficient replication of leading and lagging strand G4 structures, Related to Figure 4**

(A) Mock-depleted, DHX36-depleted, FANCJ-depleted and DHX36-FANCJ-depleted NPE supplemented with buffer, wild-type (WT) DHX36, DHX36$^{Y53A}$ (YA), or DHX36$^{E327A}$ (EA) were analyzed by Western blot with the DHX36 and FANCJ antibodies alongside a dilution series of undepleted extract.

(B) pG4$^{Lead}$ was replicated in the indicated extracts in the presence of $^{32}$P-$\alpha$-dCTP. Replication intermediates were isolated, digested with AflIII, BamHI, and ClaI (upper gel) or with AflIII and BamHI (lower gel), separated on a denaturing polyacrylamide gel, and visualized by autoradiography. The -13 to -26 products and the -1 to -3 products are indicated with a green bracket and blue bars, respectively. Defective G4 replication in the DHX36-FANCJ-depleted extracts was restored to the FANCJ single depletion level by addition of WT DHX36, but not the DHX36$^{Y53A}$ or DHX36$^{E327A}$ mutants. Thus, both ATPase and G4 recognition activities of DHX36 are required for efficient replication of the leading strand G4.

(C) Mock-depleted, DHX36-depleted, FANCJ-depleted and DHX36-FANCJ-depleted NPE supplemented with buffer, WT FANCJ, or FANCJ$^{K52R}$ (KR) were analyzed by Western blot with the DHX36 and FANCJ antibodies, alongside a dilution series of undepleted extract.

(D) pG4$^{Lead}$ was replicated in the indicated extracts in the presence of $^{32}$P-$\alpha$-dCTP. Replication products were isolated and analyzed as in (B). Defective G4 replication in the DHX36-FANCJ-depleted extracts was restored to the DHX36 single depletion level by addition of WT FANCJ, but not the FANCJ$^{K52R}$ mutant. Thus, ATPase activity of FANCJ is required for efficient replication of the leading strand G4.

(E) pG4$^{Lag}$ was replicated in the indicated extracts in the presence of $^{32}$P-$\alpha$-dCTP. Replication products were isolated and analyzed as in (B). The -13 to -26 products and the -1 to -3 products are indicated with a purple arrow and orange bars, respectively. Extension products of nascent leading and lagging strands generated after digestion with AflIII, BamHI, and ClaI are indicated on the top panels (and scheme in left panel). In the DHX36-FANCJ-depleted extracts, replication of the lagging strand was specifically compromised, suggesting that DHX36 and FANCJ play a redundant role for efficient replication of the lagging strand G4. This also shows that leading strand synthesis is not affected by a G4 on the lagging strand template. Brown hexamer, CMG; green sphere, LacI. Brown hexamer, CMG; Green sphere, LacI.

(F) pG4$^{Lag}$ was replicated and analyzed by ChIP-qPCR with DHX36 (i) and FANCJ (ii) antibodies using a primer pair for the G4 locus. Where indicated, Geminin was added to HSS to inhibit replication initiation. To compare the timing of accumulation of both proteins, DHX36 and FANCJ ChIP data in the absence of Geminin were plotted together in one graph (iii).

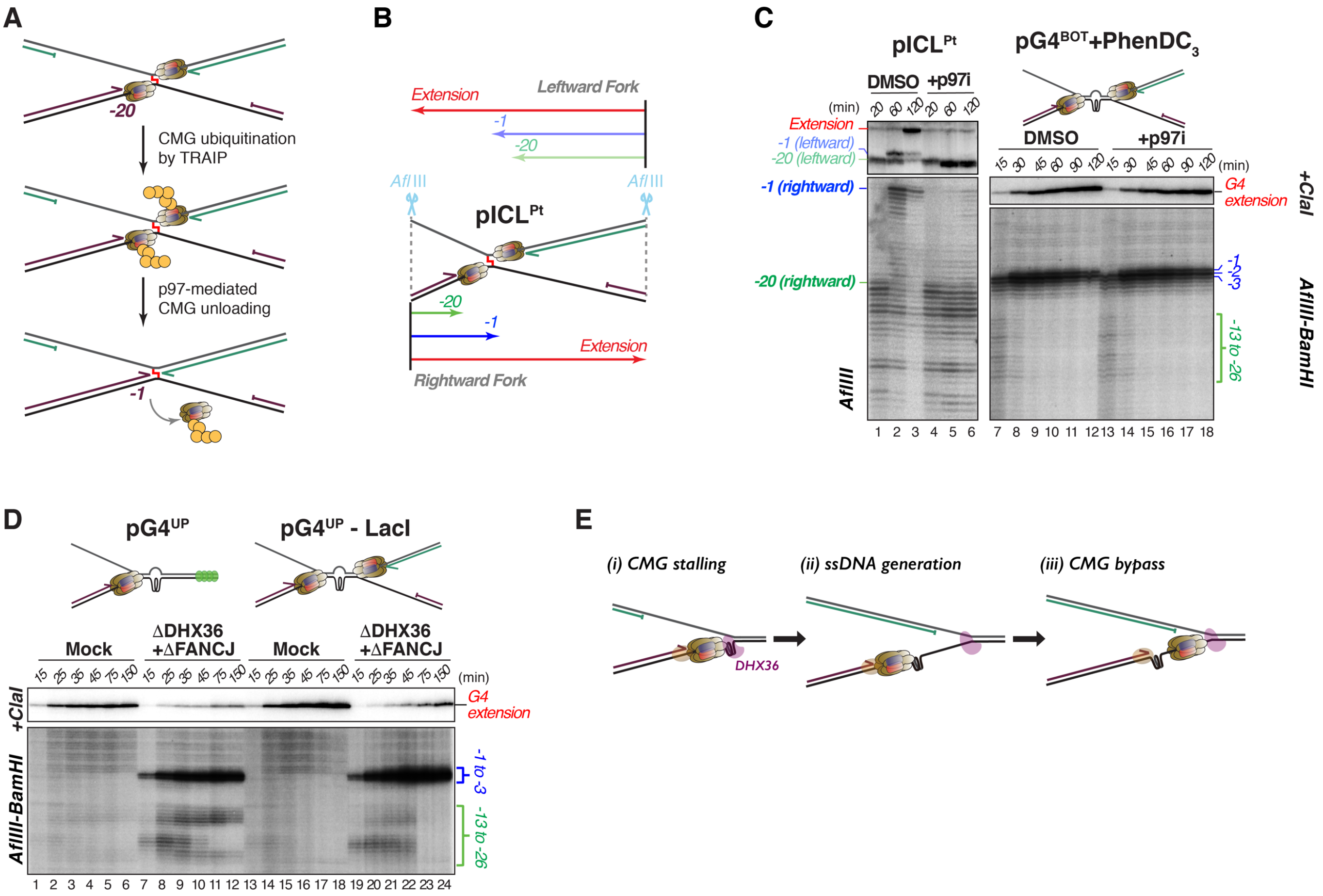

**Figure S5. DHX36 facilitates CMG bypass and not CMG unloading, Related to Figure 5**

(A) Model of CMG unloading during the repair of a cisplatin ICL. Once replication forks converge at the ICL, replication arrests at 20 to 40 nt from the ICL (CMG footprint indicated as '-20'). The ubiquitin E3 ligase TRAIP that associates with one CMG and subsequently polyubiquitinates the second CMG in trans (Wu et al., 2019) and induces p97-mediated CMG extraction from the DNA (Fullbright et al., 2016), allowing leading strand extension to 1-nt before the ICL ('-1'). Hence, dual fork convergence at an obstacle is a critical step for CMG unloading. Brown hexamer, CMG; yellow sphere, ubiquitin.

(B) Schematic representation of leading strand products of dual forks generated after replication of a plasmid that contains a site-specific cisplatin DNA interstrand crosslink ($pICL^{Pt}$) and digestion with AflIII. After approach to the -1 position, ICL unhooking, translesion synthesis, and subsequent homologous recombination result in the 685-nt extension product.

(C) $pICL^{Pt}$ and $pG4^{BOT}$ were replicated in the presence of $^{32}P$-α-dCTP with or without a p97 inhibitor NMS-873 (p97i). Replication products were isolated and digested with AflIII (lanes 1-6), AflIII, BamHI, and ClaI (lanes 7-18, upper gel) or AflIII and BamHI (lanes 7-18, lower gel), separated on a denaturing polyacrylamide gel, and visualized by autoradiography. $pG4^{BOT}$ was treated with $PhenDC_3$ prior to replication to enhance CMG stalling at the G4 structure. The -1 and -20 products generated during $pICL^{Pt}$ replication are indicated with bars on the left side. The -13 to -26 products and the -1 to -3 products generated during $pG4^{BOT}$ replication are indicated with bracket and bars on the right side, respectively. While p97i addition resulted in persistent CMG footprints at the ICL, it had no effect on $pG4^{BOT}$ replication. We thus conclude that unloading is not required for CMG to vacate the G4. This is consistent with the observation in Figure 2F that substantial MCM7 was retained on replicating $pG4^{BOT}$ at 30 minutes when most CMG vacated the G4 site (Figure S3C).

(D) $pG4^{UP}$ (G4 on bottom strand and T25 upstream of G4) was replicated in the indicated extracts in the presence of $^{32}$P-$\alpha$-dCTP. Replication intermediates were isolated and analyzed as in (C) after digestion with AflIII, BamHI, and ClaI (upper gel) or AflIII and BamHI (lower gel). Where indicated LacI was omitted. The -13 to -26 products and the -1 to -3 products are indicated with brackets. While persistent -13 to -26 products were observed in the DHX36-FANCJ-depleted extract for at least 150 minutes, these products disappeared within 75 minutes when LacI was omitted and the second fork was allowed to arrive the G4. These observations indicate that the converging replication fork facilitates CMG bypass by creating ssDNA downstream of the G4 structure.

(E) Model for CMG bypass. After CMG stalling, ssDNA is generated downstream of the G4 structure by DHX36 (I and ii). CMG subsequently bypasses the intact structure (iii).

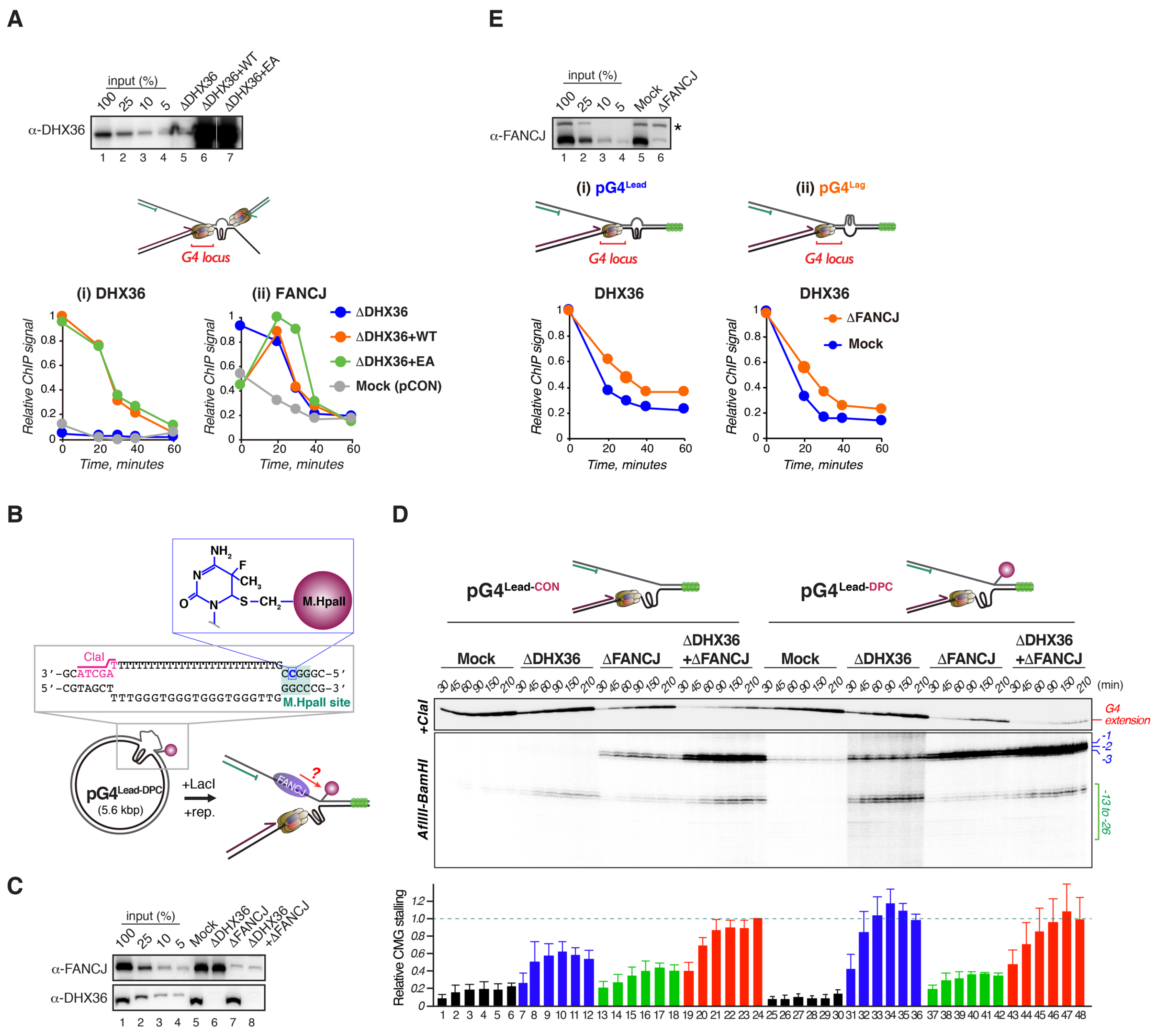

**Figure S6. DHX36 and FANCJ have partially complimentary roles when either protein is absent, Related to Figure 6**

(A) pG4$^{BOT}$ and pCON were replicated in mock-depleted or DHX36-depleted extracts supplemented with buffer, wild-type (WT) DHX36, or DHX36$^{E327A}$, and analyzed by ChIP-qPCR with DHX36 (i) and FANCJ (ii) antibodies using a primer pair for the G4 locus. The extracts used were analyzed by Western blot with the DHX36 antibody alongside a dilution series of undepleted extract. In this experiment, the G4 was replicated by dual replication forks and DHX36$^{E327A}$ dissociated from the G4 region with comparable kinetics compared to WT DHX36. Moreover, FANCJ accumulated at the G4 region in the presence of both WT DHX36 and DHX36$^{E327A}$. Therefore, the helicase switch from

DHX36 to FANCJ is not affected by the DHX36 mutation. In addition, DHX36-depletion led to robust FANCJ accumulation at the G4 locus prior to replication initiation. This initial signal likely involves FANCJ accumulation on the non-G4 template strand, as a similar accumulation was observed for a pCON control plasmid. Brown hexamer represents CMG.

(B) Schematic representation of $pG4^{Lead\text{-}DPC}$. M.HpaII (red sphere) is conjugated to the plasmid via a fluorinated cytosine (blue) at a M.HpaII recognition motif. The position of the fluorinated cytosine on the top strand is represented in the middle panel. A unique ClaI restriction site designed at a ssDNA-dsDNA junction is colored magenta. If, in the absence of DHX36, FANCJ promotes CMG bypass by translocating along the lagging strand in 3' to 5' direction, CMG bypass will be inhibited by the M.HpaII DPC in a DHX36-depleted extract in the presence of LacI (Green sphere, the bottom panel).

(C) Mock-depleted, DHX36-depleted, FANCJ-depleted, and DHX36-FANCJ-depleted NPE were analyzed by Western blot with DHX36 and FANCJ antibodies alongside a dilution series of undepleted extract.

(D) $pG4^{Lead\text{-}CON}$ and $pG4^{Lead\text{-}DPC}$ were replicated in the indicated extracts in the presence of $^{32}P$-$\alpha$-dCTP with LacI. Replication intermediates were isolated, digested with AflIII, BamHI, and ClaI (upper gel) or with AflIII and BamHI (lower gel), separated on a denaturing polyacrylamide gel, and visualized by autoradiography. The -13 to -26 products and the -1 to -3 products are indicated with a bracket and bars, respectively. The -13 to -26 products of the indicated time points were quantified and the relative intensity compared to the products in lane 24 were plotted in a bar graph with standard deviations (lower panel, n=3).

(E) $pG4^{Lead}$ and $pG4^{Lag}$ were replicated in the mock-depleted or FANCJ-depleted extract, and analyzed by ChIP-qPCR with the DHX36 antibody using a primer pair for the G4 locus. The extracts were analyzed by Western blot with the DHX36 antibody alongside a dilution series of undepleted extract (upper panel). In the FANCJ-deplete extracts, DHX36 persisted on the G4 locus of both plasmids, likely due to re-accumulation on the G4 structure. Asterisk indicates a non-specific band.

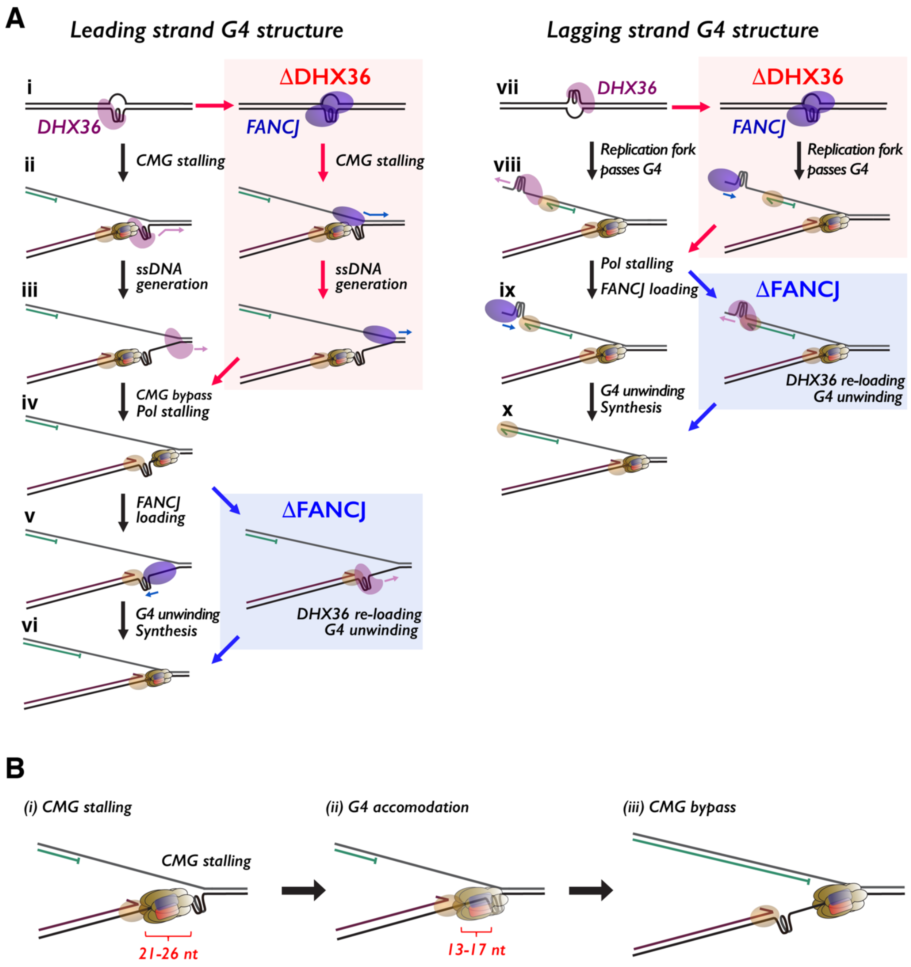


**Figure S7. Model for CMG stalling and bypass, Related to Figure 7.**

(A) Detailed models for unwinding of a leading strand G4 (left) and a lagging strand G4 (right) in unperturbed, DHX36-depleted, or FANCJ-depleted conditions. DHX36 accumulates at a G4 independently of replication (I and vii). On the leading strand template, G4 unwinding is initiated by stalling of the CMG helicase at the G4 (ii). This induces translocation of DHX36, generating ssDNA past the G4 structure (iii). The ssDNA promotes CMG bypass of the intact G4 structure, which allows leading strand extension to within a few nucleotides from the G4 (iv). This second stalling event, caused by the polymerase encountering the G4 structure, leads to the recruitment of FANCJ that unwinds the G4 (v) and allows the leading strand to extend past the G4 motif (vi). While a G4 structure on the lagging strand does not stall CMG (viii), DNA polymerase stalling is also required for the helicase switch from DHX36 to FANCJ (ix) and subsequent unwinding of the structure that allows synthesis past the motif (x). When DHX36 is absent, FANCJ accumulates at the G4 structures independently of replication and facilitates CMG bypass by translocating on the non-G4 strand (red box, left). We envision that unwinding of the lagging strand G4 in the absence of DHX36 is still DNA replication-dependent (red box, right), as FANCJ requires a free 5' end to efficiently unwind G4s (Wu et al., 2008). When FANCJ is absent, DHX36 re-accumulates at the G4s on both strands and unwinds the structures (blue boxes). Magenta oval, DHX36; blue oval, FANCJ; brown oval, DNA polymerase. (B) Detailed model for CMG bypass. First, CMG transiently stalls upon collision with a G4 structure (generating the -21 to -26 products, i). Then, it progresses at least 4 nt ahead by accommodating the structure within its N-tier channel (generating the -13 to -17 products, ii). After ssDNA is generated downstream of the G4 structure (iii), CMG bypasses the structure by opening its ring and translocating along the leading strand template strand. Given that a G4 structure is compact and the 5' and 3' end are relatively close to each other (~12 Å, see Figure S2B), both N-tier and C-tier of MCM2-7 would retain considerable contacts to ssDNA during the bypasses, avoiding accidental dissociation.
